# Integrative multiomic analysis links TDP-43-driven splicing defects to cascading proteomic disruption of ALS/FTD pathways

**DOI:** 10.1101/2025.09.29.679403

**Authors:** Velina Kozareva, Zhengde Liu, Keyana Blake, Yue A. Qi, Sasha Rollinson, Sahba Seddighi, Mohammad Alsaidi, Stanislav Tsitkov, Mercedes Prudencio, Leonard Petrucelli, Dennis W. Dickson, Anna-Leigh Brown, Pietro Fratta, Hong Joo Kim, J. Paul Taylor, Michael E. Ward, Ernest Fraenkel, Sarah E. Kargbo-Hill

## Abstract

Loss of nuclear TDP-43 is a hallmark of amyotrophic lateral sclerosis (ALS) and frontotemporal dementia (FTD). Although TDP-43 is known to regulate RNA processing, including repression of cryptic exons, we currently lack a systems-level understanding of the consequences of TDP-43 loss. To address this, we generated multiomic datasets, including RNA-seq and proteomics, from human iPSC-derived neurons depleted of TDP-43. We found that differentially spliced genes, many expressing cryptic exons, had the greatest protein reductions. Surprisingly, nearly half of differentially expressed proteins were neither mis-spliced, nor differentially expressed genes; most of these also had no reported mis-splicing in seven additional post-mortem and iPSC-derived neuron datasets. Integrative network analysis identified a high-confidence disease-specific subnetwork of over 700 interacting proteins, enriched for mRNA processing, synaptic function, and autophagy. Comparison with post-mortem ALS and FTD samples revealed convergent protein and pathway disruptions. We experimentally validated network-predicted effects of cryptic splicing in ATG4B, STMN2, and DAPK1. Our analyses reveal new TDP-43-dependent molecular cascades and nominate central genes as potential ALS/FTD therapeutic targets.

## Introduction

Amyotrophic lateral sclerosis (ALS), and frontotemporal dementia (FTD) are two devastating neurodegenerative diseases that exist on a spectrum of shared genetic and pathological characteristics. One common feature shared by nearly all ALS and about half of FTD is dysregulation of TAR DNA-binding protein-43 (TDP-43, encoded by the TARDBP gene)^1^. Pathology of TDP-43 is characterized by cytoplasmic TDP-43 accumulation and nuclear clearance of TDP-43, which can cause loss of TDP-43 nuclear functions. TDP-43 is an RNA-binding protein that regulates several aspects of RNA metabolism including transcription, translation, RNA stability^1,2^, and RNA splicing through regulation of alternative exon inclusion^3^ and polyadenylation^4,5^. Prominent among these splicing changes are those defined as “cryptic,” indicating usage of a non-canonical splice junction or intronic sequence in the mature mRNA, which can lead to a frameshift and/or premature termination^6^.

These splicing effects have been predicted and experimentally shown to affect both transcript and protein expression levels^5,7,8^. Indeed, prior studies have identified widespread mis-splicing events and global proteomic changes caused by TDP-43 depletion or dysfunction, including studies in mice^5,7,9^, neuroblastoma cell lines^10^, and induced pluripotent stem cell (iPSC)-derived neurons^8,11^. However, there has not been a systems-level investigation integrating changes across splicing, gene, and protein levels after TDP-43 loss.

Recent meta-analysis efforts have also begun to catalog mis-splicing events across ALS/FTD postmortem samples^12,13^. However, these tissue-level studies are complicated by the fact that different splicing events are observed in different cell types^7^. Overall, it remains unclear to what extent changes in protein levels following TDP-43 loss are driven by the direct role of TDP-43 in regulating splicing or other aspects of mRNA metabolism, versus being the result of secondary or further downstream effects, such as compensatory mechanisms. This knowledge is critical to understanding how TDP-43-related mis-splicing and proteomic changes contribute to neuronal degeneration in ALS-FTD and other TDP-43 proteinopathies.

To overcome these limitations, we performed multiple ‘omics studies in a unified system, investigating human iPSC-derived neurons (iNeurons) with TDP-43 depletion by CRISPR interference (CRISPRi). We examined differential splicing, RNA levels, protein levels, and TDP-43 physical interactors. We then performed integrative analysis, using a network optimization approach to generate a context specific subnetwork of protein-protein interactions (PPIs), linking splicing-related changes to global proteomic defects. These analyses revealed widespread changes in protein levels for targets of TDP-43 splicing regulation, along with wider proteomic changes for genes not directly associated with TDP-43 splicing or physical interaction. They also highlighted important processes with altered protein levels upon TDP-43 depletion, including mRNA processing, DNA-binding activity, synaptic activity, microtubule-based transport, neuron guidance, mitochondrial activity, and autophagy pathways. To investigate the disease relevance of our findings, we then analyzed several published post-mortem datasets related to TDP-43 proteinopathy. Remarkably, our optimized network, which is largely informed by iNeuron datasets, captured distinct pathways found to be disrupted across multiple ALS/FTD-related datasets, highlighting the advantage of our integrative approach. Finally, we used the network to predict downstream effects of cryptic splicing through interactions between mis-spliced genes and their protein partners. We then experimentally validated a subset of these predictions in iNeurons, including autophagy protease ATG4B and its effects on autophagy GABARAP proteins, microtubule-associated protein STMN2 and its effect on STMN1, and death-associated protein kinase DAPK1 and its effect on actin-regulator ABLIM1. Our multiomic and integrative approaches reveal novel mechanisms by which TDP-43 mis-splicing drives cascading proteomic effects related to neurodegeneration.

## Results

### Multiomic analysis reveals mis-splicing drives the strongest proteomic changes in TDP-43-depleted i^3^Neurons

To systematically examine the consequences of TDP-43 depletion on downstream gene expression, we used CRIPSRi to deplete TDP-43 from human iPSCs, which we differentiated into cortical-like glutamatergic neurons using an integrated, inducible neurogenin transcription factor cassette (hereafter i^3^Neurons, see methods)^14–16^. Since the RNA targets of TDP-43 are species- and tissue-specific, this model allows robust examination of TDP-43 function in a pure population of human neurons. We then generated multiple omics datasets from TDP-43 knockdown (KD) i^3^Neurons, including RNA-seq and mass spectrometry (MS)-based proteomics, allowing us to quantify differential gene expression, alternative and cryptic splicing, and protein expression following TDP-43 depletion (Fig 1A). We also performed affinity purification MS to identify proteins that physically interact with TDP-43. The RNA-seq data and part of the splicing analysis included here were published in our previous study focused on cryptic splicing in UNC13A^11^; we now provide more comprehensive splicing analyses of this data and report on the system-wide multiomics comparisons with additional datasets (see Methods for details).

**Figure 1:**
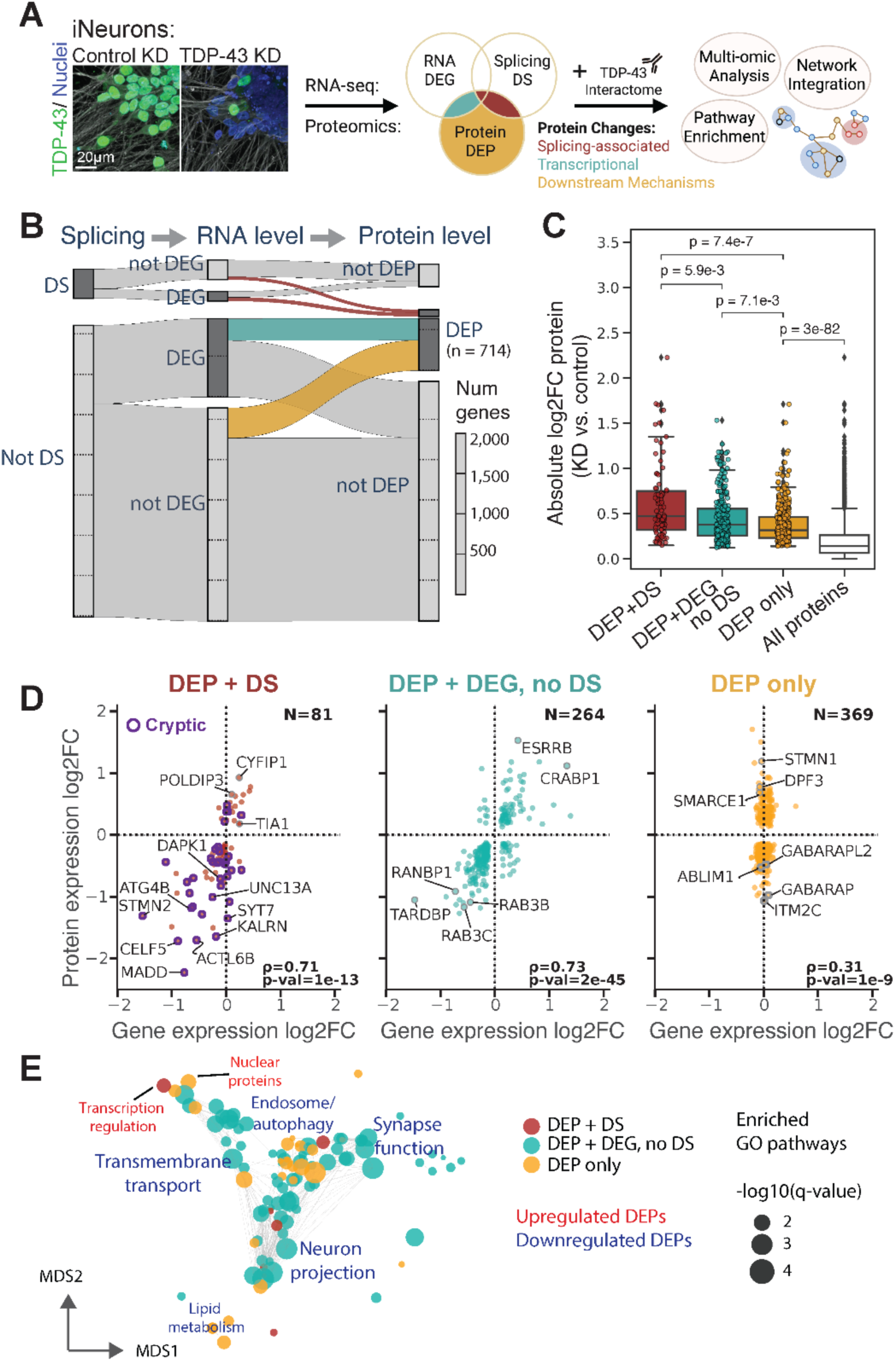
Multiomic analysis of splicing, RNA, and protein expression changes in TDP-43 KD i^3^Neurons. **A)** Schematic of data generation process and multiomic analysis strategies. Left: Confocal images of i^3^Neurons treated with a control or TDP-43-targeting guide RNA, and immunostained for TDP-43 (green). We introduce a framework used to broadly distinguish between protein level changes that can be attributed to differential splicing, transcription, or other downstream mechanisms, based on evidence of differential splicing (DS), differential gene expression (DEG), or differential protein expression (DEP). **B)** Sankey diagram quantifying genes identified as differentially spliced or expressed (color designations as in A). **C)** Distributions of absolute log2 fold changes (LFC) in protein expression across protein sets as categorized in A,B. Bonferroni-corrected p-values from two-sided Mann-Whitney-Wilcoxon tests are shown. Distribution across all proteins (including those without significant differential expression) shown in white (DEP + DS: n = 81; DEP + DEG, no DS: n = 264; DEP only: n = 369; all n = 3907). **D)** Protein expression LFC plotted against RNA expression LFC for protein sets in A,B. Proteins with cryptic splicing events outlined in purple (left). Spearman correlations and p-values indicated on each plot. N represents the number of DEPs in each category. **E)** Multi-dimensional scaling representation of GO terms found to be significantly enriched (q-value < 0.1) in overrepresentation analysis of protein sets in B,C. GO terms are grouped based on semantic similarity (Jaccard index) of member genes. Node colors represent the source set of proteins for significant enrichment. Representative summary names (text) manually derived from GO term clusters and colored by whether GO terms are enriched among up-regulated (red) or down-regulated (blue) proteins in each set of DEPs.

These datasets indicate widespread differential splicing, and transcriptomic and proteomic alterations, as expected based on previous studies from our groups and others in i^3^Neurons and other model systems (Fig S1)^2,8–10^. We were able to catalogue a combined 3,907 genes/proteins according to whether they had significant differential splicing (DS), differential RNA/gene expression (DEGs) or differential protein expression (DEPs) (Fig 1B, Methods, Table S1). This framework allowed us to broadly distinguish between TDP-43-associated protein expression effects that may be a result of altered RNA splicing (DEP+DS), altered transcript levels without splicing (DEG + DEP, no DS) and those that are most likely regulated by other downstream mechanisms with no evidence of altered RNA levels or splicing (DEP only). These downstream mechanisms may include other forms of post-transcriptional regulation or post-translational regulation, including effects from altered protein-protein interactions.

In total, we identified 714 DEPs, 81 (11%) of which had significant DS (DEP+DS), and 308 (42%) of which were also DEGs (DEG + DEP) (Fig 1B and S2A). We further characterized the DEPs as putative RNA binding targets of TDP-43, based on published individual nucleotide-resolution ultraviolet cross-linking and immunoprecipitation (iCLIP) data^17,18^ or as TDP-43 protein interactors, which we identified by Affinity-Purification Mass Spectrometry (AP-MS) experiments in iNeurons by pulling down endogenously tagged TDP-43, and supplemented by protein-protein interactome data (see Methods, Fig S2A; Table S2). Consistent with known roles of TDP-43 as an mRNA regulator, 65% of the DEPs and 70% of the DEGs had evidence of TDP-43 binding to their RNA transcripts. We also found that 83% of DEP+DS were reported RNA binding targets of TDP-43, suggesting that these DEPs are likely directly regulated by TDP-43 through its splicing function.

TDP-43 has been associated with several aspects of RNA metabolism that can impact protein levels, including splicing regulation. Interestingly, we found that genes with differential splicing had the strongest changes in protein abundance, as indicated by absolute log2 fold change, compared to all other DEPs (Fig. 1C, Fig S2A, top). To examine other potential drivers of proteomic effects, we modeled changes in protein abundance of the DEPs as a function of gene-level differential splicing, differential transcript expression, TDP-43-RNA binding and TDP-43-protein binding (see Methods). Analyzing the coefficients of this model revealed that differential splicing and differential gene expression were the strongest modifiers of protein change effect size (Fig 1C, S2B). Somewhat surprisingly, evidence of TDP-43 binding to the RNA transcript or protein had much lower effects (Fig S2B), suggesting that splice targets are particularly dependent on TDP-43 for proper protein levels compared to other RNA and protein interactors of TDP-43.

Our splicing analysis identified changes in both canonical junctions and non-canonical cryptic splicing events (see Methods). We suspected that many of the differentially expressed proteins that were also mis-spliced (DEP+DS) may undergo cryptic splicing, which can cause reduced transcript levels through nonsense-mediated decay. Consistent with this, most of the proteins in the DEP + DS set were significantly depleted, and had strong correlation with RNA expression (Spearman ⍴ = 0.71) (Fig. 1D), indicating that protein reduction in this set is generally due to decreased transcript levels. Using cryptic splicing criteria adapted from our previous analysis^11^ (see Methods), we identified 35 (43%) of these DEP+DS genes as having cryptic splicing events, including STMN2, UNC13A, ATG4B, and KALRN, with nearly all (30/35) downregulated at the protein level (Table S1). For 7 of these genes, we then confirmed cryptic exon expression and decreased gene expression using RT-qPCR (Fig. S3). The remaining non-cryptic alternatively spliced genes were more likely to be associated with increased protein abundance (Fig 1D), including CYFIP1 (LFC=0.9), an actin regulator, which has been reported upregulated in FTD brains^19^, the RNA binding protein, POLDIP3 (LFC=0.66), previously reported in ALS tissue^20^ and TIA1 (LFC=0.18), a stress-granule protein with mutations causative for ALS^21^.

Overall, these analyses highlight that TDP-43 loss drives significant protein-level changes in neurons through regulation of alternative and cryptic splicing, with cryptic splicing predominantly leading to protein reduction, consistent with previous studies of TDP-43-mediated splicing repression^4^.

### Differentially expressed, but not mis-spliced, proteins disrupt distinct disease-related pathways after TDP-43 loss

While mis-splicing drives the strongest proteomic changes following TDP-43 depletion, most DEPs (633, 89%) did not have associated differential splicing, but may still impact disease-related pathways. Some of these proteins were also differentially expressed at the transcript level (DEP + DEG, no DS, Fig 1D, center) and displayed high protein/RNA fold-change correlation (Spearman ⍴ = 0.73). Genes in this category, especially those with strong concordance of transcript and protein changes, may reflect direct, non-splicing-mediated targets of TDP-43 regulation or indirect transcriptional responses to TDP-43 loss. We also characterized protein-level changes that had no accompanying transcript-level change (DEP only). These had the smallest effect sizes overall, but comprised nearly half of all differentially abundant proteins (n = 369) (Fig 1C,D). These changes may be due to a combination of post-transcriptional regulation, such as defects in RNA transport, which is a known function of TDP-43^22,23^, post-translational loss of stabilizing protein interactions, or other mechanisms.

We next asked whether pathway-level proteomic disruptions were primarily driven by mis-splicing or if there were significant impacts from other non-mis-spliced genes. Gene ontology (GO)^24,25^ term enrichment analysis on the DEP groups (DEP+DS, DEP+DEG no DS, DEP only) revealed both shared and distinct clusters of pathways associated with these protein sets (Fig 1E, Fig S2C, Table S3). In particular, we found enrichment of neuron-specific processes (synaptic function, neuron projection) across all three sets of proteins. This is consistent with a model where mis-splicing of key synaptic genes (UNC13A, SYT7, KALRN) may cause cascading effects on other synaptic proteins potentially through both transcriptional and post-transcriptional mechanisms. However, some pathways only emerged as significantly impacted among the protein-specific changes (DEP only). These include processes associated with autophagy and lipid metabolism (Fig 1E, Fig S2C). Further analysis of DEPs with discordant RNA expression (see Methods) revealed additional pathway-level patterns including relatively increased protein abundance in chromatin remodeling and mRNA splicing, and decreased protein abundance in ribosomal, plasma membrane, and organelle assembly pathways (Fig S2D, Table S4). These DEP only dominated effects may represent secondary cascades where mis-regulation of a few genes (or even a single gene) affects downstream protein targets. For example, cryptic splicing and loss of the autophagy gene ATG4B can cause post-translational modifications on the autophagy gene ATG3^26^, and may also explain other autophagy-related changes in the DEP only group, such as reduced levels of GABARAP and GABARAPL2 (Fig 1D).

### A compendium of mis-splicing events associated with TDP-43 proteinopathy or TDP-43 depletion

To corroborate the gene-level mis-splicing we characterized in i^3^Neurons, and to gain a more comprehensive view of mis-splicing events related to TDP-43 proteinopathy, we built a compendium of TDP-43-associated mis-splicing events from published studies. We compiled sets of genes with reported differential or cryptic splicing from seven additional analyses, including an independent experiment in i^3^Neurons with TDP-43 depletion^27^, several analyses of post-mortem ALS/FTD tissues^11,28,29^, a recent meta-analysis of *in vitro* and post-mortem datasets^12^, and two recent studies of cryptic splicing in i^3^Neurons with TDP-43 depletion and NMD inhibition^30,31^ (Table S5, Methods).

Overall, these analyses largely differed in their reported sets of differentially spliced genes even between i^3^Neuron studies (Fig. S4A), with the majority of genes (3265/4240) being detected as DS or cryptic in only a single analysis (Fig. S4B, Table S6). However, our set of 1352 DS genes (see Methods) derived from our i^3^Neuron data showed substantial overlap with other sets, with 585 DS genes reported as mis-spliced in at least one other analysis (Fig. S4A). As suggested in other studies^12^, we expect that the discrepancy between these reported gene sets may be due to differences in cell-type/tissue composition, context-specific effects, or analysis methods with variable inclusion of low probability events. Nevertheless, there is a small set of genes (13) that were robustly detected as mis-spliced (appearing in ≥ 6 analyses) (Fig. S4B). These include well-studied genes like STMN2, UNC13A, and KALRN, along with others like PTPRT and PTPRD, two members of the tyrosine phosphatase receptor family that have not been directly linked to ALS/FTD, but are involved in synaptic function and have been associated with neurodevelopmental and neurodegenerative disorders^32,33^. Additionally, 653 genes were detected as mis-spliced in both i^3^Neuron and post-mortem contexts, and these more robustly detected genes are associated with higher magnitude protein-level changes in our i^3^Neuron dataset (Fig. S4C,D). An important caveat of this compendium meta-analysis is variable criteria between studies for defining particular splicing events as “cryptic.” Some genes meet cryptic criteria in some analyses but not others (for example, CYFIP1 and TIA1). We therefore consider differential splicing on a gene level more broadly, rather than focusing specifically on “cryptic” events or junctions, in many of our following analyses.

We also examined three analyses of cryptic or alternative polyadenylation (C/APA)^34,35^ (Table S5), as these mechanisms of mRNA regulation have also recently been shown to impact expression levels of TDP-43 targets^35–37^. These analyses identified a combined set of 2355 genes with C/APA, although as with mis-splicing datasets, there was limited overlap between gene lists (Fig. S4E, Table S6). Comparing these to the mis-spliced genes, 754 genes had both reported DS and C/APA, though these genes were not associated with more extreme protein-level changes than other DS or C/APA genes (Fig. S4F).

This compendium implicates additional genes not detected in our original differential splicing analysis, as either mis-spliced or with alternative polyadenylation. We observe significant overlap between these additional genes and our i^3^Neuron DEPs (257/714), suggesting these DEPs may be partially driven by TDP-43-associated mis-splicing or alternative polyadenylation (Fig. S4G). While some of these DEPs may not actually undergo significant alternative splicing or polyadenylation in our experimental system, we highlight prominent expression changes in genes such as RAP1GAP, ARMC10, and PDE1B (Fig. S4G) that are potentially explained by these mechanisms. However, the majority of DEPs (376/714) still show no association with alternative splicing or polyadenylation in any dataset examined, consistent with our model that primary TDP-43 targets drive secondary cascading proteomic effects.

### Differential TF activity contributes to transcriptional regulation after TDP-43 loss

Our analyses identified 3877 differentially expressed genes (DEGs) following TDP-43 depletion (including those without protein-level changes), 90% of which were not associated with differential splicing detected in our model system. We reasoned that some of these gene expression changes may be caused by effects on other gene regulators, such as transcription factors (TFs) or RNA-binding proteins (RBPs), particularly for TFs or RBPs known to be direct targets of TDP-43 regulation (Fig. 2A). Thus, to investigate potentially relevant activity of TFs we used a database of 1186 TF regulons (collecTRI^38^) to infer differential TF activity from our differential gene expression data (Methods). We identified 34 TFs with significant differential activity (DA) (FDR-adjusted p-value < 0.1), with 29 of these inferred to be more active in TDP-43 depleted i^3^Neurons (Fig. 2B, Fig. S5A, Table S7). Fifteen of these DA TFs were also differentially expressed on a gene level, with 10/15 showing concordance between direction of inferred activity and expression. Overall, we also observed a modest but significant positive correlation between inferred TF activity and gene expression (Spearman ⍴ = 0.2, p = 4e-5, Fig 2B). TFs were sparsely detected in our proteomics data, which may explain why associations between DEPs and inferred activity did not reach statistical significance.

**Figure 2:**
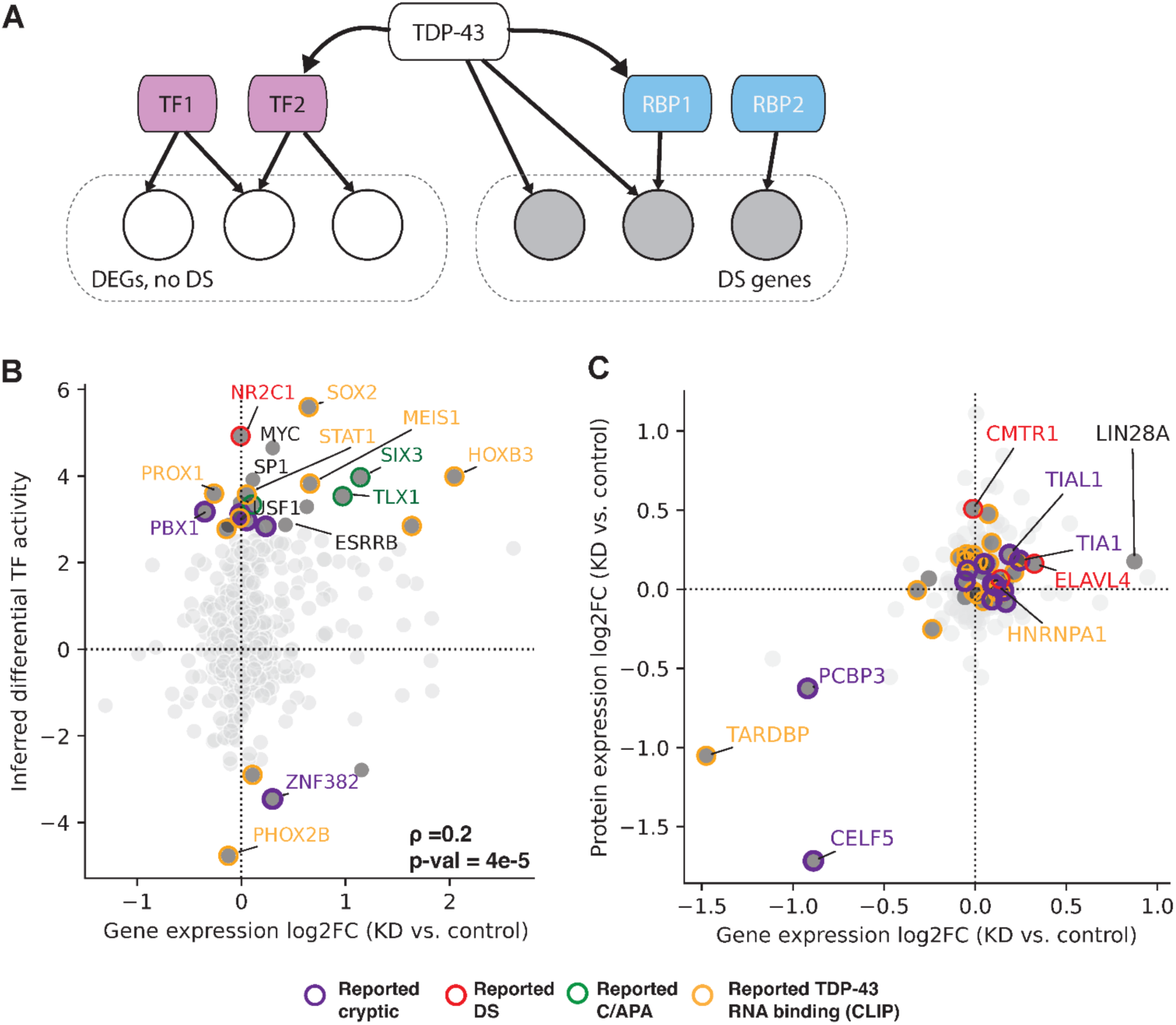
Inferred differential activity of TFs and RBPs. **A)** Simplified model of potential regulatory interactions with TFs and RBPs modulating differential gene expression and splicing downstream of TDP-43. **B)** Inferred TF activity level plotted against RNA LFC for corresponding gene (all 423 TFs quantified in i^3^Neuron RNA-seq data). **C)** Protein expression LFC plotted against RNA expression LFC for all 221 RBPs quantified in i^3^Neuron RNAseq/proteomics. TFs with significant differential activity or RBPs with significant motif enrichment (considered DA, see Methods) in dark grey and annotated with borders indicating evidence of DS, C/APA, or TDP-43 RNA binding based on this and other studies. Select TF/RBPs labeled.

Of the inferred active TFs, 21 had evidence of direct regulation by TDP-43 either through differential splicing or RNA-binding, including 8/10 of the most significant inferred TFs (Fig 2B, S5A). Several of these TFs have been previously associated with TDP-43 dysfunction. In particular, SIX3 and TLX1 were recently reported to undergo 3’Ext cryptic polyadenylation in TDP-43 depleted i^3^Neurons, leading to increased RNA and translation levels^34^. In addition, PHOX2B, a putative RNA target of TDP-43 by CLIP^18^ and inferred here to have significantly decreased TF activity, has been shown to regulate neurite length and axon resilience, with reduced transcript stability when TDP-43 is mutated^39^. SP1, which is inferred more active, has previously been shown to regulate p11 (S100A10), which is neuroprotective in SOD1 models of ALS^40^. In addition, several of our inferred TFs were also identified as part of an integrative transcriptional-metabolic analysis across 6 neurodegenerative diseases, including USF1, SP1, ESRRB, MYC, KLF4, STAT1, and PBX1^41^, suggesting some of these responses may be consistent with broader neurodegenerative signatures.

To determine if TFs identified by this analysis are potentially responsible for regulation of disease associated pathways following TDP-43 loss, we examined the putative TF targets and compared them to differentially expressed genes and proteins without differential splicing (DEG+DEP, no DS). We found that 22% of these DEG+DEPs were putative TF targets; among all DEGs (including those without quantified protein levels) 23% were putative targets. Target genes regulated by DA TFs with positive activity were enriched for processes related to ribosomal function, neuronal migration, and regulation of TF activity, while those regulated by putatively less active TFs were related to forebrain development, neuron regeneration, and energy homeostasis (Fig S5B, Table S8). In fact, many inferred TFs seem to co-regulate each other (Fig S5C), providing additional evidence that TDP-43 loss disrupts complex regulatory networks in neurons.

### RNA-binding protein (RBP) involvement in direct and indirect splicing effects after TDP-43 loss

Given the prominent role of RBPs in ALS/FTD pathophysiology^42,43^, we hypothesized that RBPs might be differentially regulated upon TDP-43 loss and contribute to indirect effects on target genes (Fig. 2A). We identified 27 RBPs that were differentially expressed in our TDP-43 KD i^3^Neurons at the protein level, the majority with increased expression. These include several known ALS genes, such as ELAVL3 (decreased), and TIA1 (increased). We also found many RBPs (155), that were physical interactors of TDP-43, including known ALS genes, MATR3, TAF15, HNRNPA1, and HNRNPA2B1 (Table S2)^44^.

To determine if these or other RBPs might contribute to some of the differential splicing events observed upon TDP-43 KD, we tested differentially spliced exon-exon junctions for enrichment of known RBP binding motifs^45^ (for 175 RBPs, see Methods). We performed stratified analyses using subsets of DS junctions to identify RBPs that may be acting in conjunction (or in competition) with TDP-43, along with those that may contribute to indirect differential splicing effects. Briefly, we first generated an inclusive subset of all exon-level mis-splicing events (DS Set 1), and progressively excluded junctions with reported TDP-43 CLIP binding sites (DS Set 2) and nearby TDP-43 motifs (DS Set 3) to identify splicing events that may be due to indirect regulation by other RBPs (see Methods, Fig S6A).

In total, we identified significant enrichment of 62 unique motifs, associated with 53 RBPs (dark gray dots in Fig 2C, Table S9, Methods), which we consider to be inferred differentially active RBPs. As expected, our analysis showed highly significant enrichment for UG-rich motifs (associated with TDP-43 binding^18^) in the broadest subset (DS Set 1; Fig S6B,C). Accordingly, the smaller DS subsets (DS Sets 2/3) with less evidence of direct TDP-43 binding showed less significant enrichment of this and other TDP-43 associated motifs^46^ (Fig S6C). Conversely, several RBP-motif pairings displayed more significant enrichment in the narrower subsets, including PCBP3 and LIN28A (Fig S7A, downregulated panel), suggesting these may regulate non-TDP-43 splicing events.

Among the RBPs differentially expressed on a protein level (|LFC| > 0.5), we found a moderately significant enrichment of inferred RBPs (p = 0.03, Fisher’s exact test). Furthermore, most of the inferred RBPs (44/53) themselves show evidence of either missplicing or regulation by TDP-43 (Fig 2C). Examples include cryptic splice targets such as CELF5, and other DS genes with decreased protein levels like PCBP3. In addition, this analysis highlighted several RBPs previously associated with ALS/FTD, including HNRNPA1^47^ and TIA1^48^. Interestingly, a prior study also identified similar enrichment of TIA1 CLIP binding sites near TDP-43 associated cryptic exons^49^. Additionally, we were able to identify motif enrichment patterns consistent with known binding profiles for these RBPs based on published CLIP datasets (TIA1^50^, HNRNPA1^51^) (Fig. S7B,C).

Overall, these motif enrichments are suggestive of potential RBP binding near junctions with differential splicing after TDP-43 loss, but it is possible this binding does not contribute directly to splicing effects. Further investigation of genome-wide binding profiles and differential splicing targets is needed to determine the functional significance of less well studied inferred RBPs.

### Integrative network analysis reveals mis-spliced genes at the heart of proteomic disruptions

To gain a systems-level understanding of the specific pathways altered by TDP-43 loss in neurons, and to investigate the relationships between mis-spliced genes and broader proteomic changes, we generated a context-specific subnetwork of protein-protein interactions using the Prize Collecting Steiner Forest (PCSF) algorithm^52^ (Methods). This network approach integrates multiple sources of input nodes (genes and proteins) with a set of known protein-protein and protein-metabolite interactions (PPMIs), generating a pruned network that retains high-confidence interactions between nodes of interest while filtering out poorly connected nodes. This method has previously been used to identify significant interactions and functional groups across multiple disease contexts^53–57^.

Our subnetwork was constructed using input nodes derived from the DEPs, inferred TFs and RBPs, and other DS genes not detected in the proteomics (see Methods, Table S10). The final network consists of 723 nodes (Fig 3A, Table S11), and includes 175 “Steiner” nodes, proteins that are not included in the original input sets, but that are predicted to interact significantly with other input nodes (Methods). These may represent important functional partners of proteins impacted by TDP-43 loss that were not detected in our experimental datasets.

**Figure 3:**
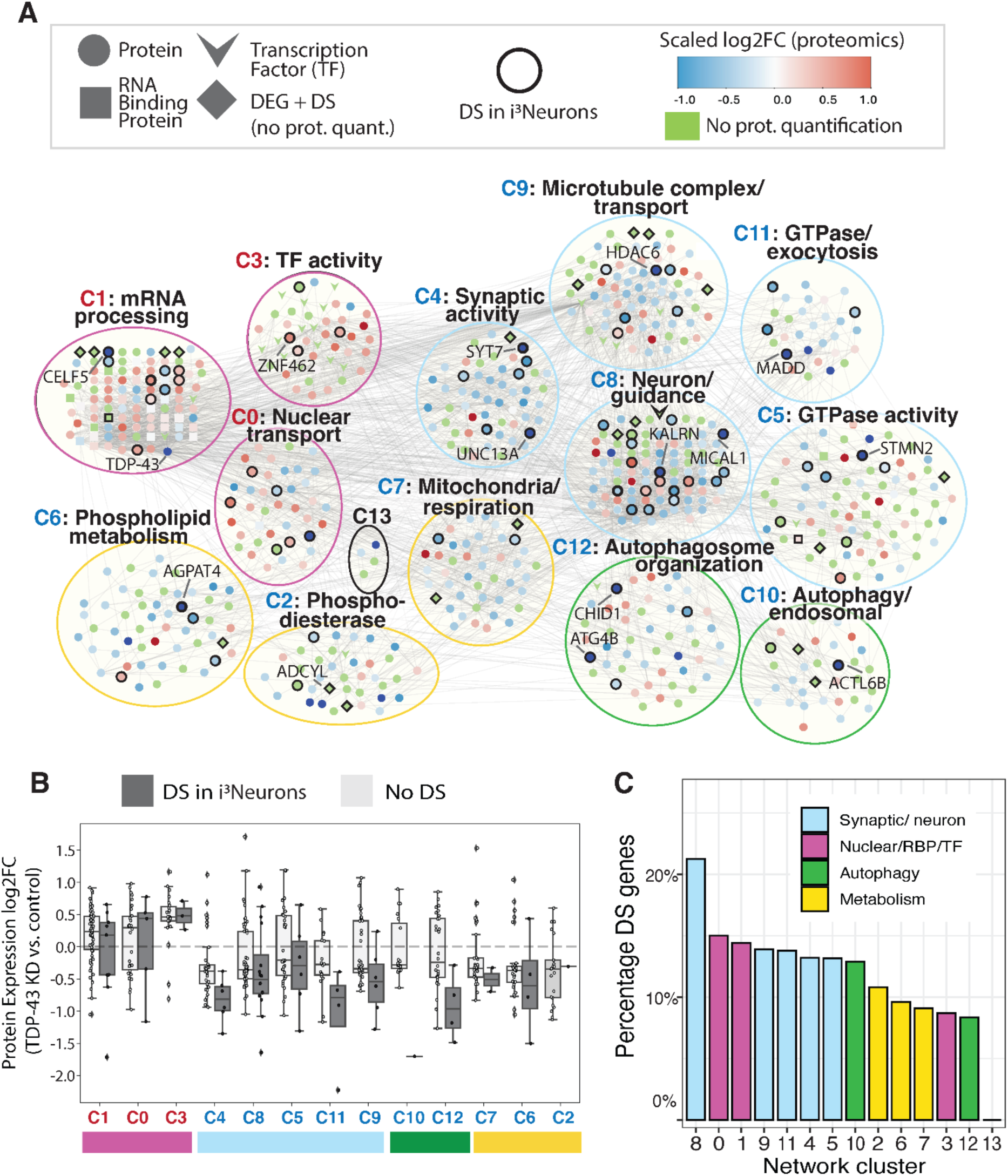
Protein-protein interaction subnetwork reveals links between mis-spliced genes and wider proteomic changes. **A)** Overview of clustered TDP-43-specific PPI network, highlighting genes with differential splicing in our i^3^Neuron dataset, and summarized pathway enrichment for each network cluster. Clusters are organized by “meta-clusters” based on similar functional pathway enrichments. The network contains 723 nodes consisting of proteins (including RBPs and TFs) and metabolites (Methods). **B)** Boxplot showing distributions of protein expression LFC values for each network cluster, labeled as in A, separated by DS status. **C)** Barplot indicating proportion of nodes in network clusters with DS. Colors indicate “meta-clusters” of network clusters grouped by functional similarity of enriched GO terms.

To better characterize functional groups of proteins within the network, we applied Leiden community detection and performed GO term enrichment on the resulting clusters (C0-C12; Fig 3A, Table S12). This revealed enrichment of several sets of related processes (Fig 3A), including neuron-related pathways like synaptic function (C4), neuron projection (C8), and microtubule transport (C9), and nuclear-localized functions like mRNA processing (C1) and TF activity (C3). Other cluster groups were enriched for autophagy (C12, C10) and metabolic processes, including mitochondrial activity (C7), and phospholipid metabolism (C6).

Although most clusters consisted of both up- and downregulated proteins, the majority of functional groups showed a strong bias towards decreased protein expression (blue labels; Fig 3A, B). However, clusters related to mRNA processing and splicing (C1), TF activity (C3), and nuclear transport (C0) showed primarily increased expression (red labels). This upregulation of proteins related to transcriptional and splicing activity could suggest a compensatory response to loss of normal RNA regulation, driven by reduction of TDP-43 and dysfunction of other RBPs themselves altered by TDP-43 depletion.

Notably, genes with differential splicing were distributed throughout nearly every cluster, indicating their potential involvement in a wide array of disrupted processes (Fig 3A,C). Several clusters had particularly high proportions of DS proteins, including the neuron projection (C8) and mRNA processing (C1) clusters (Fig 3C). Cluster C8 (a subset of which is highlighted in Fig S8) includes several known TDP-43 splice targets related to neuronal projection and cytoskeletal/actin organization, including KALRN and MYO18A^31^. Other DS proteins include genes like MICAL1, DAPK1, and CYFIP1, which have been previously linked with neurological disorders including epilepsy^58^, Alzheimer’s disease^59^, and other dementias^60^. Interestingly, although most C8 cluster proteins show decreased expression, CYFIP1 is one of several increased actin regulators, suggesting that dysfunction of this pathway is driven by both reduction and increase of key regulators.

In the mRNA processing cluster (C1), we find TDP-43/TARDBP itself, along with many of its direct protein interactors (40/96 nodes in the cluster). Also within this cluster are many proteins known to affect RNA splicing, including several hnRNPs (HNRNPH3, HNRNPCL1, HNRNPH1, HNRNPA1, HNRNPAB, HNRNPH2), and Serine/Arginine-Rich Splicing Factors (SRSF10, SRSF1, SRSF2), and a core component of the U1 small nuclear ribonucleoprotein, SNRPA (Fig S8).

The main synaptic cluster (C4), contains not only previously reported cryptic exon-containing genes, UNC13A, SYT7, DNM1, and CADPS, but also many other down-regulated synaptic genes: Synaptotagmins (SYT1, SYT2, SYT5), synapsin family members (SYN1, SYN2), and synaptophysin (SYP) (Fig S8). While prior work from our groups and others points to a significant role for the UNC13A cryptic exon, this analysis highlights additional downregulation of synaptic machinery, which may be downstream of UNC13A or act in parallel.

Closely related to both cluster C4 and cluster C8 is a cluster enriched for microtubule-related transport and dynein activity (Cluster C9). Many proteins within this cluster (DCTN1/2, DCTN4, DYNLL2) show significantly depleted protein levels (Fig S8), consistent with impaired transport in neurons after TDP-43 loss of function. Similar microtubule and transport defects have been reported in previous studies of TDP-43 dysfunction, including in recent work profiling protein expression in patients with TDP-43-related dementias^61^. Importantly, mutations in DCTN1 have been linked to ALS, FTD and Perry disease^62–64^, and lead to increased aggregation of TDP-43. DCTN1 dysfunction as well as general neuronal trafficking defects have been implicated in TDP-43 aggregation, potentially through direct interaction or disruption of stress granule dynamics^64^.

As a resource for the community, we have also created an interactive, annotated version of the network available at https://fraenkel-lab.github.io/tdp43-multiomic-network/network (Data and Code Availability).

### Proteomics analysis of C9-FTLD postmortem patient brain reveals differential expression in mis-spliced genes

To investigate the extent to which our i^3^Neuron datasets model effects present in patient brains, we analyzed frontal cortices of C9-FTLD-TDP patients and healthy controls (Fig. 4A), which we previously generated as a benchmark dataset for our proteomics pipeline^65^. These samples (N=10 patients, N=10 healthy controls, Table S13) were selected for severe TDP-43 pathology, as measured by high phospho-TDP-43 levels from a larger cohort^66^. By focusing on severe TDP-43 pathology, we are able to identify TDP-43-dependent pathological protein changes for co-analysis with TDP-43 KD i^3^Neurons. From this postmortem cohort, we observe 579 up-regulated and 913 down-regulated proteins (Fig. 4B), with strong concordance between our postmortem and i^3^Neuron datasets (128 of 191 shared DEPs) (Fig. 4C). We also observe that many of the DEPs detected in the post-mortem dataset were differentially spliced in i^3^Neurons with TDP-43 depletion (141/1492) (Fig. 4B), and that down-regulated, concordant proteins were especially enriched for differential splicing in the i^3^Neurons (26/104 DEPs) (Fig. 4C). Pathway analysis of the C9-FTLD dataset revealed both overlapping signatures with i^3^Neurons and distinct patterns. For example, similar enriched GO terms indicate decreased expression of Golgi vesicle transport, synaptic activity, macroautophagy, and GTPase activity pathways. However, additional terms, such as increased levels of immune response and extracellular matrix proteins, may reflect non-neuronal contributions to disease (Fig. S9A). While some of these DEPs in post-mortem samples may arise from non-splicing or non-neuronal contributions, these results are consistent with mis-splicing contributing significantly to protein loss in patients with FTLD-TDP pathology.

**Figure 4:**
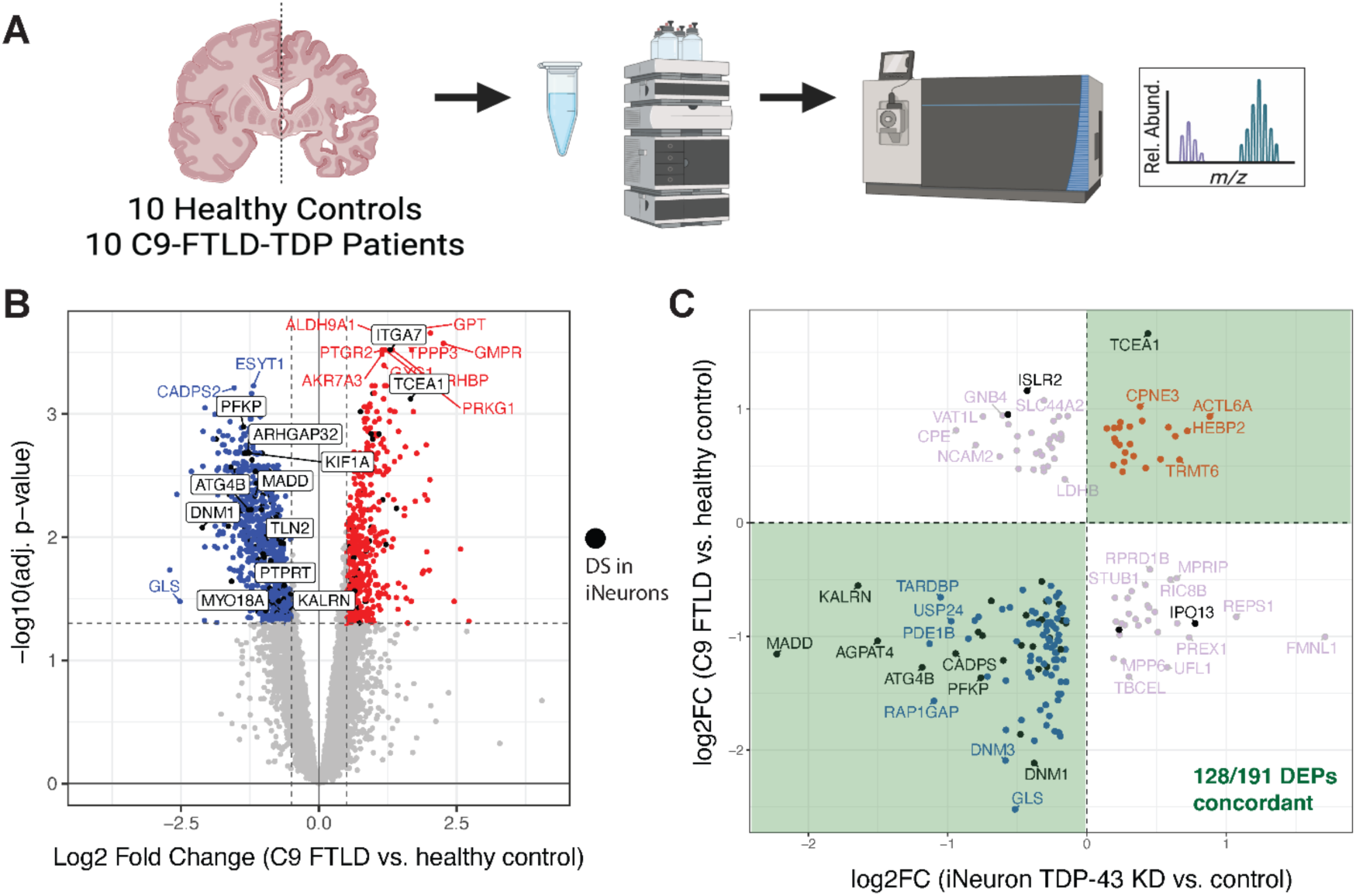
Differential analysis of bulk proteomics from C9-FTLD-TDP brains. **A)** Schematic of mass spectrometry-based proteomics experiment on C9-FTLD-TDP patient brains and healthy controls. **B)** Volcano plot showing differentially expressed proteins between C9-FTLD and healthy control samples. Dot color indicates whether the protein is significantly differentially expressed (non-grey: FDR < 0.05, |log2FC| > 0.5). Black dots and boxed gene labels indicate genes with differential splicing in i^3^Neurons with TDP-43 KD (our dataset). Only notable or previously studied proteins are labeled. **C)** Scatter plot comparing LFC expression values in post-mortem C9-FTLD vs. controls and i^3^Neurons TDP-43 KD vs. controls, for 191 proteins that were DE in both analyses. Black points indicate genes that also have differential splicing in i^3^Neurons with TDP-43 KD (our dataset).

### Network highlights pathways disrupted across multiple post-mortem ALS/FTD contexts

To more broadly determine whether the pathways uncovered in our network, primarily driven by i^3^Neuron datasets, reflect disease biology in patients, we surveyed a panel of four additional published proteomic and transcriptomic studies of ALS/FTD spanning multiple tissues (Table S14). These datasets include bulk proteomics from frontal cortex of FTD brains^67^; single-cell proteomics of ALS spinal-cord motor neurons stratified by TDP-43 inclusion status^68^; bulk RNAseq from FACS-sorted neuronal nuclei from frontal cortex of ALS-FTD cases^69^; and bulk RNA-seq from a range of central nervous system tissues generated by the NYGC ALS consortium^11,70^.

We processed each dataset using standardized methods to obtain differential expression signatures (Table S15), followed by separate GO enrichment analyses for genes or proteins increased and decreased in disease (Table S16). When we compared the resulting pathway signatures with the functional groups defined by our network, we found that prominent enriched pathways from nearly all clusters were represented in at least one post-mortem context (Fig. 5A). Notably, several pathways that were not significantly enriched when considering either i^3^Neuron RNA or protein changes alone (e.g., autophagy and mitochondrial activity) emerged as disease-relevant through this integrative comparison, and many of the associated enrichments involved genes that are mis-spliced and present within the network.

**Figure 5:**
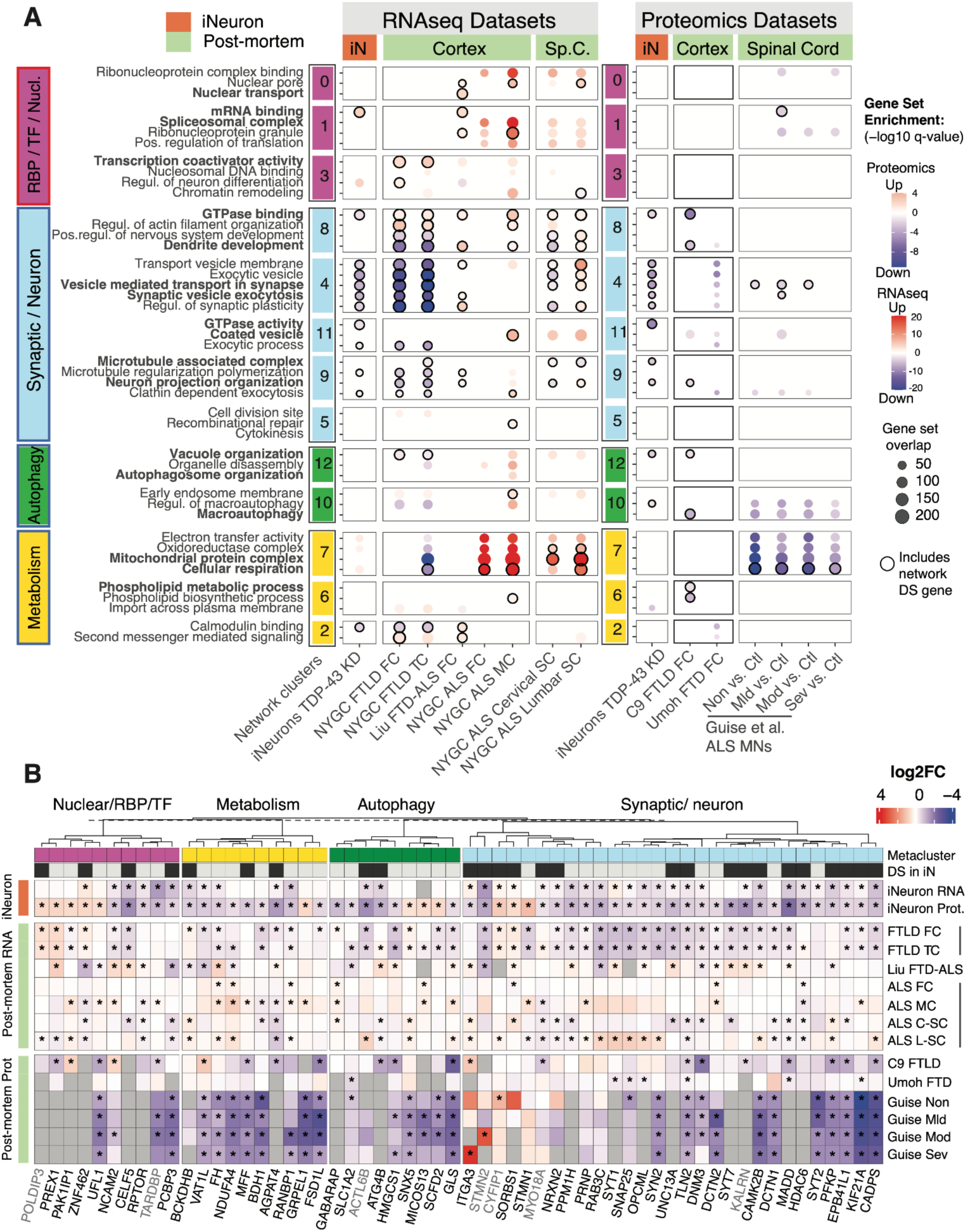
Integrated analysis captures diverse pathways and genes disrupted across multiple post-mortem disease contexts. **A)** Dotplot of i^3^Neuron and post-mortem patient datasets for MSigDB GO terms enriched across the integrated network, organized by the modality of dataset (RNAseq, proteomics). (See Methods, Table S16 for all enriched GO terms). Cluster pathway enrichments organized by metacluster and functional similarity (left). Highly significant terms are bolded. Dot colors indicate whether GO terms are enriched among upregulated or downregulated genes, with saturation corresponding to -log10(q-value). Black border on dot indicates one or more DS gene(s) from the network appear in the enriched GO term. **B)** Heatmap of gene and protein expression LFC values across post-mortem patient datasets for select DEPs from our network, separated by metaclusters as defined in Figure 2. Hierarchical clustering of genes using LFC values is performed within each metacluster separately. Network cluster membership and DS status indicated for each gene. Most displayed genes were found to have significant DE in multiple datasets (see Methods for criteria); additional genes of interest not meeting strict inclusion criteria are shown in grey text. * BH-adjusted p-value < 0.05.

Pathways related to synaptic function and microtubule transport (network clusters 4 and 9) showed largely consistent down-regulation across multiple datasets, whereas autophagy (clusters 10, 12) and mitochondrial processes (cluster 7) displayed context-dependent directional changes. Such differences likely reflect variation in cell-type composition and possibly distinct disease mechanisms across regions; for instance, spinal-cord bulk samples contain fewer neurons than cortical tissues^11,71^. Consistent with this, the single-cell spinal cord proteomics dataset (consisting of motor neurons) did exhibit synaptic and exocytosis deficits, while the bulk spinal cord RNA-seq showed more variable changes across these and other pathways. Additionally, pathway profiles in FTLD cortex aligned most closely with our i^3^Neuron network, echoing recent work showing that TDP-43-associated splicing signatures in vitro resemble those seen in FTLD cortex more than in ALS cortex or spinal cord^12^.

Beyond pathway-level agreement, we observed significant expression changes in many individual network genes and proteins in patient tissue (Fig. 5B). For example, the differentially spliced cytoskeletal and synaptic regulators TLN2^72^ and EPB41L1 (Cluster C4) were consistently decreased across nearly all post-mortem datasets, suggesting they might serve as versatile indicators of TDP-43 proteinopathy. Conversely, CYFIP1, highlighted earlier in the actin cytoskeleton-related cluster C8, was elevated across most contexts, also consistent with the i^3^Neuron results (Fig. 5B). Other mis-spliced genes, such as ZNF462 and ITGA3, displayed divergent expression patterns, likely reflecting regional differences in cell type composition, splicing, variable TDP-43 pathology, or additional regulatory inputs.

Finally, the disease relevance of potential regulators predicted from our i^3^Neuron analysis was also supported by post-mortem data. This includes several transcription factors with inferred activity changes, multiple RBPs, and Steiner nodes, which were often differentially expressed in concordance with our predictions (Fig. S9B). Together, these results underscore that the pathways and proteins highlighted by our network analysis capture core aspects of ALS/FTD biology while also revealing context-specific variations that may illuminate tissue- or disease-subtype heterogeneity.

### Network analysis identifies secondary consequences of TDP-43 splicing regulation

We anticipated that protein loss of TDP-43 splice targets may have downstream effects on other interacting proteins and related pathways. Our integrative network, by robust prioritization of significant protein-protein interactions, can be used to predict specific functional interactions between direct TDP-43 targets (“primary targets”) and putative candidates for post-transcriptional or post-translational regulation (DEPs only) (“secondary targets”).

To determine if loss of primary targets alone would be sufficient to alter secondary targets, we depleted candidate primary targets using CRIPSRi in i^3^Neurons and performed western blots to look at levels of secondary targets for comparison with TDP-43 KD i^3^Neurons. We reasoned that if a secondary target was driven by mis-splicing and loss of a primary target, then loss of that primary target should phenocopy loss of TDP-43 for protein levels of the secondary target. To select candidates for testing, we focused on secondary DEPs with network neighbors that were mis-spliced and downregulated (DS+DEP). We also focused on secondary candidates with no evidence of differential gene expression or splicing (Fig. S10A-S10C), eliminating those likely to be driven by independent mechanisms. Candidates were further selected based on protein change effect sizes, degree of discordance between RNA/protein LFC (Fig. S10D).

We tested 6 candidate secondary targets (GABARAP, GABARAPL2, ITM2C, STMN1, ABLIM1, and BAIAP2), connected to 3 primary targets (ATG4B, STMN2, and DAPK1) (Fig 6A-D). We observed that 4 secondary targets were independently driven by knockdown of the primary target (Fig. 6E-J). These included GABARAP and GABARAPL2, which are interacting partners of ATG4B (primary, with a cryptic exon), which appears in the autophagosome enriched cluster (Fig. 3A, Cluster 12). Prior studies established a strong connection between dysfunction of autophagy and ALS/FTD^73,74^. ATG4B is a cysteine protease that cleaves LC3 and GABARAP family proteins during autophagosome formation and maturation. A previous study identified ATG4B cryptic splicing in ALS tissues and its role in regulating LC3B-protein conjugation.^26^ However, this is the first study to implicate altered GABARAP and GABARAPL2 protein levels downstream of ATG4B cryptic splicing. In our analysis of post-mortem datasets, we observed significantly decreased protein expression of GABARAPL2 in spinal cord motor neurons (Fig. 6K), though not in cortical proteomics datasets. Interestingly, both GABARAP and GABARAPL2 had elevated transcript levels across multiple bulk RNAseq datasets, including spinal cord (Fig. 6K), potentially indicating tissue-specific effects or a compensatory transcriptional response.

**Figure 6:**
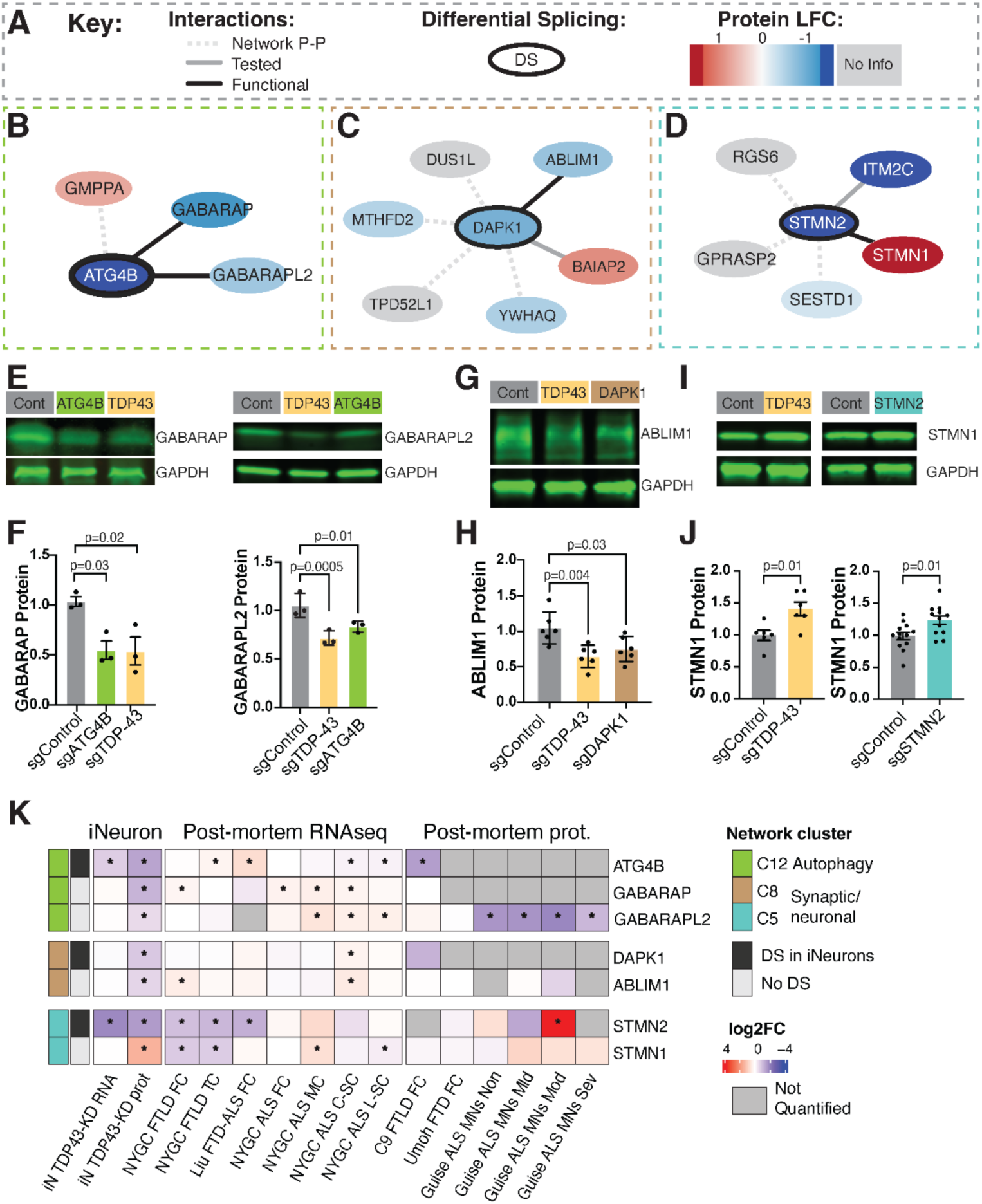
Predicted functional relationships between TDP-43 splice targets and secondary protein effects, experimentally validated. **A-D)** Schematic showing primary splice targets and cluster-specific protein-protein interaction partners with key (A), ATG4B (B), DAPK1 (C), and STMN2 (D). **E)** Western blots of GABARAP (left) and GABARAPL2 (right) showing protein levels for representative Control KD, ATG4B KD and TDP-43 KD i^3^Neurons. GAPDH is used as a loading control. **F)** Quantification of E for N=3 biological replicates per group. Statistics using ordinary one-way ANOVA with Dunnett’s multiple comparisons test between indicated groups. Error bars show SEM. **G)** Western blot of ABLIM1 showing protein levels for representative Control KD, TDP-43 KD and DAPK1 KD i^3^Neurons. **H)** Quantification of G for N=6 biological replicates per group. Statistics using ordinary one-way ANOVA with Dunnett’s multiple comparisons test between indicated groups. Error bars show SEM. **I)** Western blots of STMN1 showing protein levels for representative Control KD and TDP-43 KD (left) and Control KD and STMN2 KD (right). **J)** Quantification of I for N=6 biological replicates per group (left), and for N=12 or 13 biological replicates per group (right). Statistics using Welch’s t test; Error bars show SEM. **K)** Heatmap comparing gene and protein expression LFC values across post-mortem patient datasets for functionally tested genes/proteins. * BH-adjusted p-value < 0.05.

We also identified ABLIM1 as the secondary target of DAPK1. A previous study identified ABLIM1 as one of 600 DAPK1 interactors in cortical neurons^75^. ABLIM1 is reduced upon depletion of either TDP-43 or the primary TDP-43 target, DAPK1 (Fig. 6G,H). ABLIM1 is an actin-binding protein involved in axon outgrowth and guidance^76^. While ABLIM1 has been connected to ALS as a splice target of FUS^77^, contributions to neurodegeneration are unknown. DAPK1 or Death Associated Protein Kinase 1 is a stress-responsive protein that is upregulated in Alzheimer’s disease, and stroke^78^. DAPK1 has also recently been linked to a SOD1 model of ALS where it appears elevated and contributes to motor neuron death^79^. Since up-regulation of DAPK1 signals apoptosis, lowered expression due to mis-splicing may actually function as a neuroprotective response to TDP-43 reduction.

Due to its prominent role as a cryptic exon, we also tested if STMN2 acts as a primary target to potential secondary target STMN1. Interestingly we observed that STMN1 levels were higher in neurons with either TDP-43 KD or STMN2 KD (Fig. 6I-J; S10E, S10F). We speculate that STMN1 levels may increase to compensate for decreased STMN2. Similar increases in STMN1 levels were observed in several post-mortem ALS studies (Fig. 6K), suggesting this may also occur in disease contexts. While we observed decreased levels of BAIAP2 and ITM2C following TDP-43 KD, as predicted by our proteomics, we did not see any changes upon DAPK1 KD or STMN2 KD respectively (Fig. S10G, S10H).

These studies highlight the utility of the integrative network to predict potential cascading relationships among targets, where TDP-43-induced mis-splicing can drive secondary protein effects, including in pathways known to be associated with disease. However, our CRISPRi studies only establish that primary targets ATG4B, DAPK1 and STMN2 are capable of regulating proteomic changes in their secondary targets. Individual mechanistic studies will be needed to determine if these secondary targets are driven entirely by or only partially by loss of their corresponding primary target.

## Discussion

TDP-43 loss of function in neurons represents an early stage of disease pathogenesis^80,81^, causing widespread defects in RNA metabolism that may then trigger downstream events resulting in neuronal dysfunction, death, and disease progression. Prior transcriptomic and proteomic studies have collectively identified thousands of TDP-43-dependent transcript and protein level changes^2,8–10,12^. However, with so many putative effectors, it remains a major challenge to identify which events stem directly from TDP-43 loss, and which events may be secondary or tertiary consequences of a pathological cascade.

To address these challenges, here we explore the cellular consequences of TDP-43 loss with integrative analyses of splicing, transcriptomic, and proteomic datasets in a single neuronal cell type, iPSC-derived glutamatergic neurons or i^3^Neurons. We stratify differentially spliced genes, differentially expressed genes and proteins, TDP-43-protein interactions, and TDP-43-RNA interactions^17,18^, and observe that the strongest protein-level effects are associated with mis-spliced genes, but that the majority of differentially expressed proteins are not associated with differential splicing or gene-level changes (Figure 1). While our analyses highlight the critical role of TDP-43 in splicing and gene level regulation, they also prompted us to explore how alteration of primary targets may cause secondary consequences on other proteins. Thus, we computationally investigated putative primary targets including transcription factors and RBPs that themselves may regulate many downstream targets (Figure 2). This revealed several potential transcriptional regulators that are involved in disease-relevant neuronal pathways.

To gain a systems-level view of the state of the neuron upon TDP-43 loss, we also generated a high-confidence disease-specific subnetwork of protein-protein interactions, integrating our i^3^Neuron datasets and other neuron-centric data sources (Figure 3). Importantly, this network represents the first of its kind, revealing cascading effects of TDP-43 loss and highlighting relevant interactions between TDP-43 targets and other proteins, which we present to the community as an interactive resource (link: https://fraenkel-lab.github.io/tdp43-multiomic-network/network).

We then compared our new network, which was largely informed by i^3^Neuron datasets, with our recent FTLD post-mortem proteomic dataset. The FTLD samples showed significant disruption of several genes and pathways also highlighted by our network and multiomic results (Figure 4). We then expanded our analyses to include other previously published ALS and FTD postmortem RNA-seq and proteomic datasets. This wider cross-comparison helped to reveal additional patterns of pathway and gene dysfunction across multiple disease contexts (Figure 5). Finally, we experimentally validated network-predicted functional relationships between cryptic-exon-expressing genes and their protein interactors (Figure 6). These combined bioinformatic and experimental approaches support a model of cascading effects where TDP-43 acts on primary RNA targets either through regulating splicing, RNA stabilization or another mechanism. These primary targets are then effectors, which regulate additional targets that become altered upon TDP-43 loss of function.

Our integrative network and multiomic analyses provide a new framework for deciphering TDP-43-associated pathways. Many of the pathways identified in our network have been previously found to be associated with TDP-43 proteinopathy, including synaptic processes, neuron outgrowth, autophagy, and RBP disruption. However our combination of transcriptomic and proteomic analyses, including protein-protein interactions, provides additional mechanistic insights into these pathway-level effects, and suggests new hypotheses to explain protein-level effects.

For example, while most neuron-associated clusters were down-regulated, interestingly the mRNA processing cluster appeared to be up-regulated (Figure 3). We speculate that this increase in expression for RBPs and splicing factors may be a compensatory mechanism in response to TDP-43 loss. This model is supported by our multiomics, RBP activity inference, and network clustering analyses. Intriguingly, we also observed up-regulation of similar pathways in multiple post-mortem RNAseq datasets, suggesting that this is potentially a transcriptionally driven effect that persists even in later stages of disease. While this up-regulation is not observed across all post-mortem proteomic analyses, a concurrent proteomic study of hippocampal tissue from patients with Limbic-predominant Age-related TDP-43 Encephalopathy Neuropathologic Change (LATE-NC) and Alzheimer’s Disease,^61^ identified similar increases in mRNA processing and splicing proteins, highlighting another context where this up-regulation occurs possibly as a compensatory response to TDP-43 loss.

Another disease-associated pathway that appeared across several of our analyses is autophagy. We observed significant down-regulation of autophagy proteins in our multiomic and network analyses, but many of these proteins, with the exception of cryptic-exon-containing gene ATG4B, did not have corresponding RNA-level changes. We suspected that loss of ATG4B may drive a molecular cascade of proteomic changes in other autophagy proteins. We showed that ATG4B depletion was sufficient to decrease GABARAP and GABARAPL2 protein levels similar to TDP-43 KD. In addition to autophagy, our network analyses identified several disease-relevant pathways, including lipid metabolism, mitochondrial activity, and trafficking. Additional studies will be needed in the future to investigate the putative molecular cascades of these pathways.

We note several important limitations of these datasets that will need to be addressed in future studies. First, our data was generated from i^3^Neurons with TDP-43 KD, which fail to capture many aspects of disease, including contributions by mature synaptic networks, non-neuronal cells, and effects of aging. We also focus on TDP-43 loss of function, while gain-of-function mechanisms, including those that may contribute to splicing dysregulation,^82^ are not captured in our i^3^Neuron data. We may capture some of this signature in our analyses of patient datasets, however additional studies using models of TDP-43 aggregation will be needed to differentiate gain-of-function and loss-of-function mechanisms of TDP-43 dysfunction. Lastly, quantification of protein and RNA expression is an imperfect estimation of functional protein and transcript levels, particularly for proteomics where many impacted proteins may be below levels of detection. This is especially relevant for lowly-expressed proteins such as transcription factors, and future experiments could use more targeted proteomics approaches and functional assays to better quantify protein levels and activity.

Nonetheless, together these analyses illustrate the core role of TDP-43 in splicing regulation, and identify particularly disruptive splicing events associated with strong protein loss, which are robustly identified across i^3^Neuron and patient datasets. Through network integration and functional validation experiments, we provide support for a cascading model of TDP-43-dependent effects. Understanding the specific effectors of this cascade is critical to identifying effective target(s) for therapeutic intervention, and predicting possible secondary effects of modulating individual targets.

## Methods

### iPSC culture and iNeuron differentiation

Most iPSCs used in this study were from the WTC11 line, which was derived from a healthy male donor and obtained from the Coriell Cell Repository. The WTC11 cell line has a normal male karyotype and carries no genetic mutations associated with ALS–FTD risk. APMS experiments were performed using the BJFF6 iPSC line, derived from a healthy male donor, and modified to endogenously tag TARDBP with eGFP, as described in the next section.

iPSC culture maintenance was performed following previously published protocols.^15^ Briefly, tissue culture (TC)-treated petri dishes were coated with human embryonic stem cells (hESC)-qualified Matrigel (Corning, 354277) for a minimum of 30 minutes before plating iPSCs. iPSCs were then maintained in Essential 8 medium (Thermo Fisher, A1517001). Passaging of iPSCs was performed using either Accutase (Thermo Fisher, A1110501) or 0.5 M EDTA in PBS. Cultures were supplemented with 10 μM ROCK inhibitor (Chroman 1; MedChemExpress, HY-15392) on the day of splitting and thawing to promote iPSC survival. Culture media was changed daily or every other day.

To generate iPSC-derived neurons, we used i^3^Neurons, in which WTC11 was previously genetically modified to insert a doxycycline-inducible mouse neurogenin-2 expression cassette into the AAVS1 safe harbor site.^14^ To start neuronal differentiation, 2-4 millions of accutase-dissociated iPSCs were plated onto matrigel-coated 10cm TC petri dish (0.5–1 million iPSCs per well of a 6-well plate and 10–20 million iPSCs for a 15-cm dish) at Day 0 in differentiation media, consisting of KnockOut DMEM/F-12 medium (Thermo Fisher, 12660012) supplemented with 1X N-2 supplement (Thermo Fisher, 17502048), 1X GlutaMAX supplement (Thermo Fisher, 35050061), 1X MEM nonessential amino acids (Thermo Fisher, 11140050), 10 μM ROCK inhibitor (Chroman 1; MedChemExpress, HY-15392), and 2 μg/ml doxycycline (Sigma-Aldrich, D9891). Media were changed daily during differentiation.

On Day 3, cells were individualized with accutase and subcultured onto TC-treated 6-well plate or 15cm petri dish coated with poly-l-ornithine (PLO; Alamanda Polymers, 27378-49-0) in BrainPhys neuronal culture medium (STEMCELL Technologies, 05790) supplemented with 1× B27 Plus Supplement (Thermo Fisher, A3582801), 10 ng/ml NT-3 recombinant protein (Thermo Fisher, 450-03), 10 ng/ml BDNF recombinant protein (Thermo Fisher, 450-02), 1000 ng/mL mouse laminin (Thermo Fisher, 23017015), and 2 μg/ml doxycycline (Sigma-Aldrich, D9891). To feed i^3^Neuron, half media changes with warm BrainPhys neuronal culture medium were performed three times a week. 2 million i^3^Neuron were plated per well of a 6-well plate, and 10 million for 10cm plate. Unless otherwise noted, i^3^Neurons were collected at Day 17 (14 days post-subculture onto PLO-coated plates) for subsequent experiments.

### Endogenous tagging of the TARDBP/TDP-43 gene

The *TDP43* eGFP tagged BJFF.6 iPSC line was generated using CRISPR-Cas9 technology. Briefly, BJFF.6 iPSCs (RRID:CVCL_VU02) were pretreated with mTeSR (Stem Cell Technologies) supplemented with 1X RevitaCell (Thermo Fisher Scientific) for 1 hour. Then, approximately 1X10^6^ cells were transiently co-transfected with 1.15ug of SM32.hTDP43.g9 plasmid (cloned into Addgene #43860; spacer below), 1.5ug of p3s-Cas9HC plasmid (Addgene #43945), 2.8ug of SM32.eGFP donor (sequence below), and 500ng of pCE-mp53DD (Addgene #41856). The transfection was performed via nucleofection (Lonza, 4D-Nucleofector™ X-unit) using solution P3 and program CA-137 in a small (20ul) cuvette according to the manufacturer’s recommended protocol. Five days post-transfection cells were sorted on viability and plated onto Vitronectin XF (Stem Cell Technologies) coated plates into prewarmed (37C) mTeSR media supplemented with 1X CloneR (Stem Cell Technologies). Clones were expanded, screened, and verified for the desired knockin via PCR assays and sequencing. Editing construct sequences and relevant primers are listed in Table 1 below.

**Table 1:**
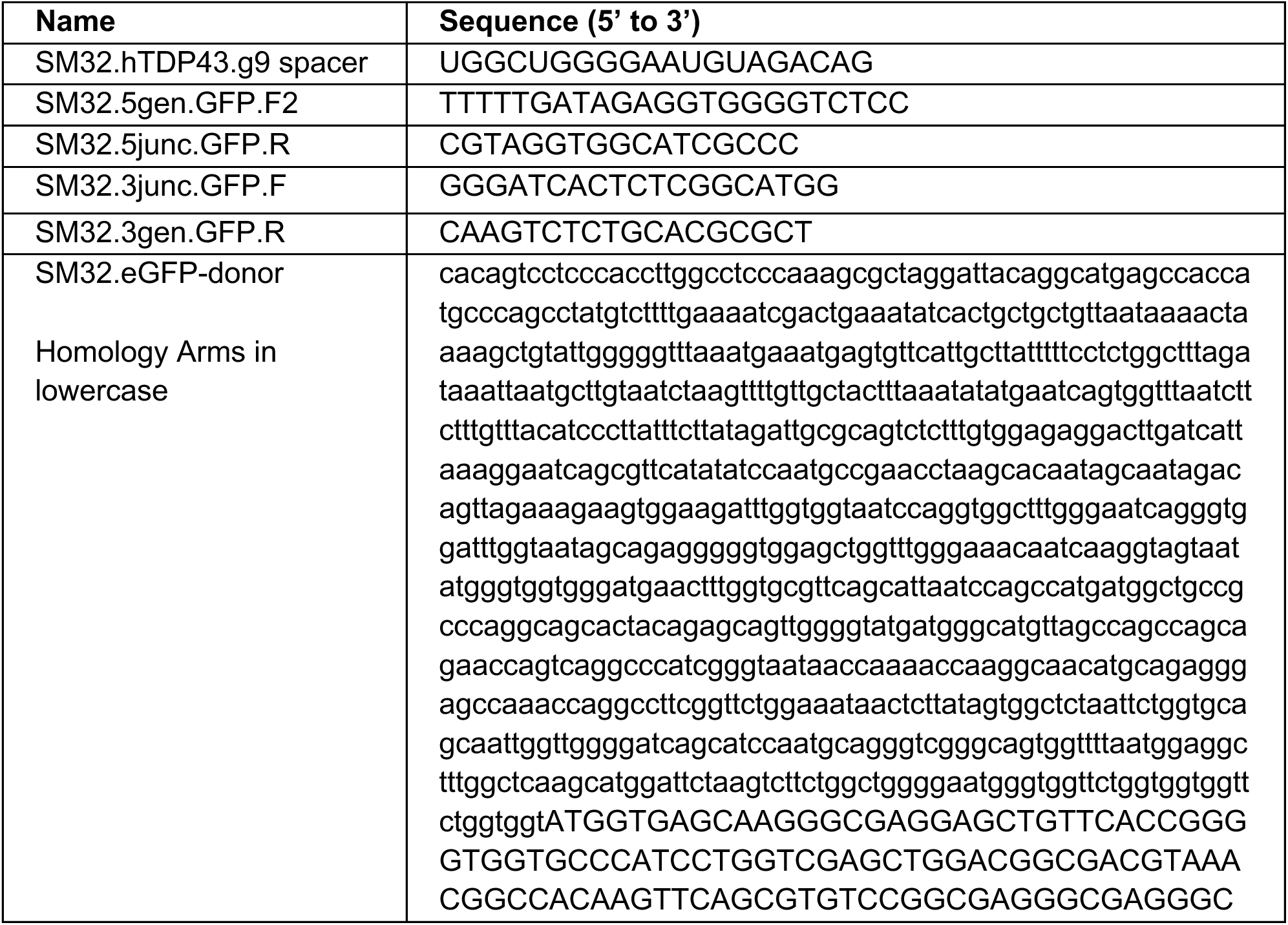
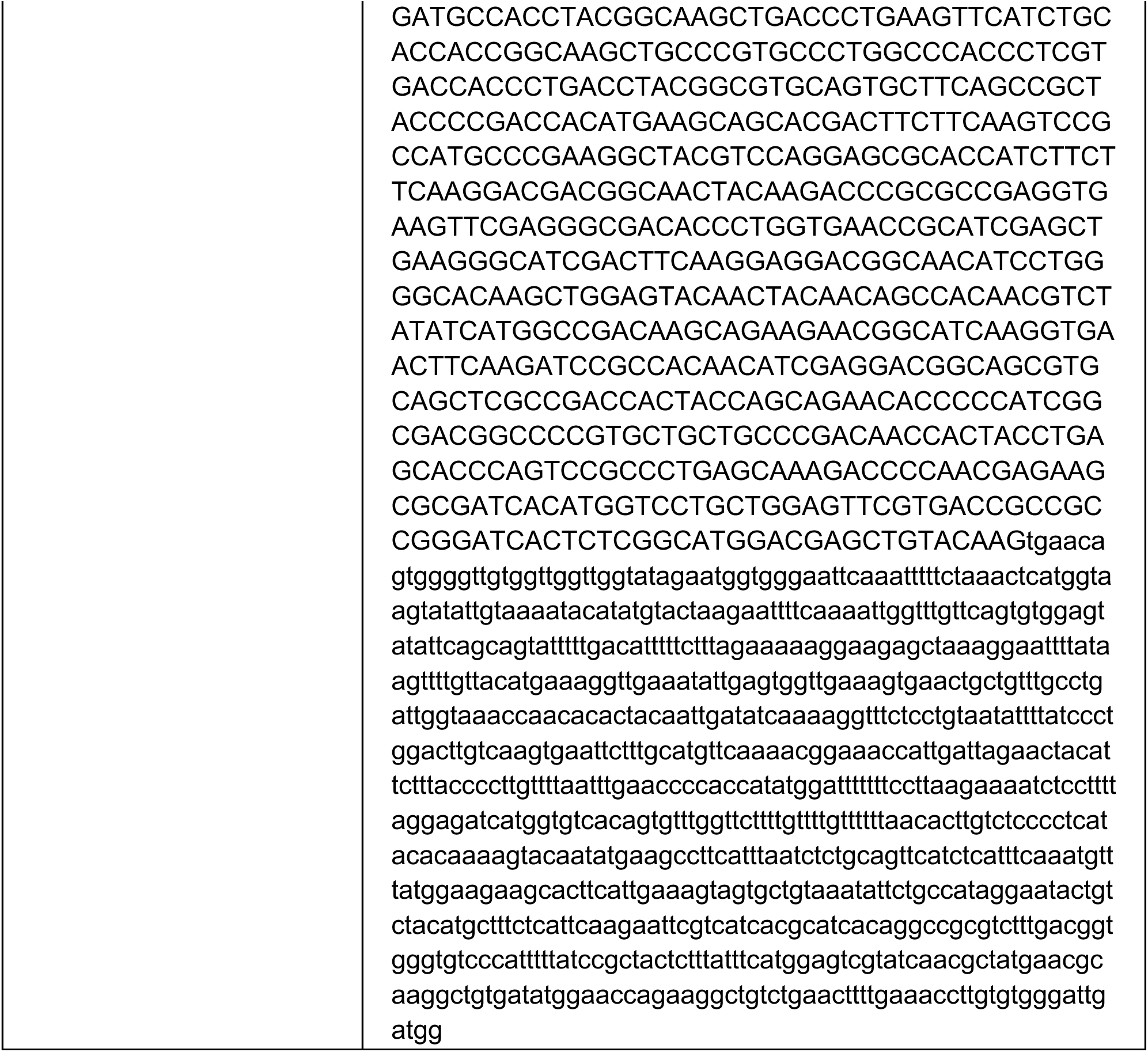

This line was differentiated into i^3^Neurons using the same differentiation protocol as above. Briefly, a doxycycline-inducible NGN2 construct was delivered to the iPSCs using Piggybac integration. The piggybac plasmid and a plasmid expressing EF1alpha-transposase were transfected into iPSCs using Lipofectamine Stem Reagent (Thermo Fisher Scientific), following the manufacturer’s protocol. Transfected iPSCs were then selected for with hygromycin and iPSCs were differentiated using doxycycline induction.

### CRISPRi Knockdown

The WTC11 iPS cells used in this study were engineered to express a CAG-driven, enzymatically dead Cas9 (dCas9-BFP-KRAB) cassette integrated into the CLYBL safe harbor locus.^16^ To accomplish knockdown, lentivirus was used to deliver sgRNA targeting the corresponding gene or a non-targeting control guide to iPSC as described below. Accutase-individualized iPS cells in suspension were treated with viruses and then plated onto Matrigel-coated TC petri dish in E8 media supplemented with ROCK inhibitor. PBS wash and media change with fresh E8 or E8 + ROCK inhibitor on lentivirus-transduced cells were performed the next morning. One or two days after media change, iPSCs were replated onto a new marigel-coated TC plate in E8 media with ROCK inhibitor and 10 μg/ml puromycin (Sigma-Aldrich, P4512) for overnight selection. Cells were then expanded for one to two days before initiating neuronal differentiation. Knockdown efficiency was tested with RT-qPCR or western blot on Day 3 or Day 17 i^3^Neuron.

The sgRNA sequence were cloned into pU6-sgRNA EF1Alpha-puro-T2A-BFP vector (Addgene #60955). The following guide sequences were used: non-targeting control (GTCCACCCTTATCTAGGCTA), TDP-43 (GGGAAGTCAGCCGTGAGACC), ATG4B (GCTGCGGAAAGACCGACCCC), STMN2 (GAAGGGTCCGGCTACAGCAG), DAPK1 (GGAGGCTGCTTCGGAGTGTG).

### Lentiviral production

Lenti-X Human Embryonic Kidney 293T (HEK293T) cells were used to produce lentivirus. Lenti-X HEK293T cells were plated onto poly-L-ornithine (PLO)-coated TC-treated petri dish at a density of 3 millions cells per well of a 6-well plate, 15 millions cells per 10-cm dish, or 45 millions cells per 15cm dish in warm DMEM, high glucose, GlutaMAX Supplement media (Thermo Fisher, 10566024) supplemented with 10% Fetal Bovine Serum (FBS) (Thermo Fisher, A5670701; Sigma-Aldrich, F2442) and then culture overnight. On the next morning, cells were transfected with Lipofectamine 3000 (Thermo Fisher, L3000015), P3000, psPAX2, pMD2G, pAdVantage, and the lenti-vector of interest in room temperature Opti-MEM (Thermo Fisher, 31985062). Media changes were performed the next morning with warm DMEM, high glucose, GlutaMAX Supplement media + FBS supplemented with ViralBoost (Alstem, VB100). Post-transfection cells were maintained for 2 to 3 days without changing media. Then, the media were collected and cell debris were removed by centrifugation. The lentivirus were 1:10 concentrated in PBS from the supernatant by using the Lenti-X concentrator (Takara, 631231). Concentrated lentivirus were aliquoted and stored in −80 °C.

### RT-qPCR

i^3^Neurons were collected on day 14 post dox-induction with TRI Reagent (Zymo Research R2050-1-200). RNA was isolated following the Direct-zol™ RNA MiniPrep Plus Kit (Zymo Research R2072) protocol. A nanodrop was used to measure RNA concentrations. RNA was normalized and 300 ng was used to reverse transcribe cDNA using the High-Capacity RNA-to-cDNA™ Kit (ThermoFisher 4387406) protocol. qPCR was used to analyze gene expression with SsoAdvanced Universal SYBR Green Supermix (BIO-RAD 1725271), primer (see Table 2) concentration of 500 nM and either undiluted or 1:10 diluted cDNA. Samples were run on a CFX Connect Real-Time PCR Detection System (BIO RAD) and quantified using the ΔΔ*C*t method.

**Table 2:**
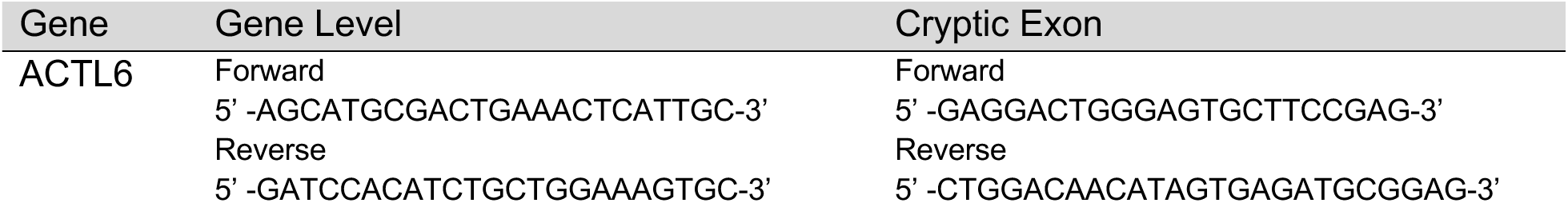
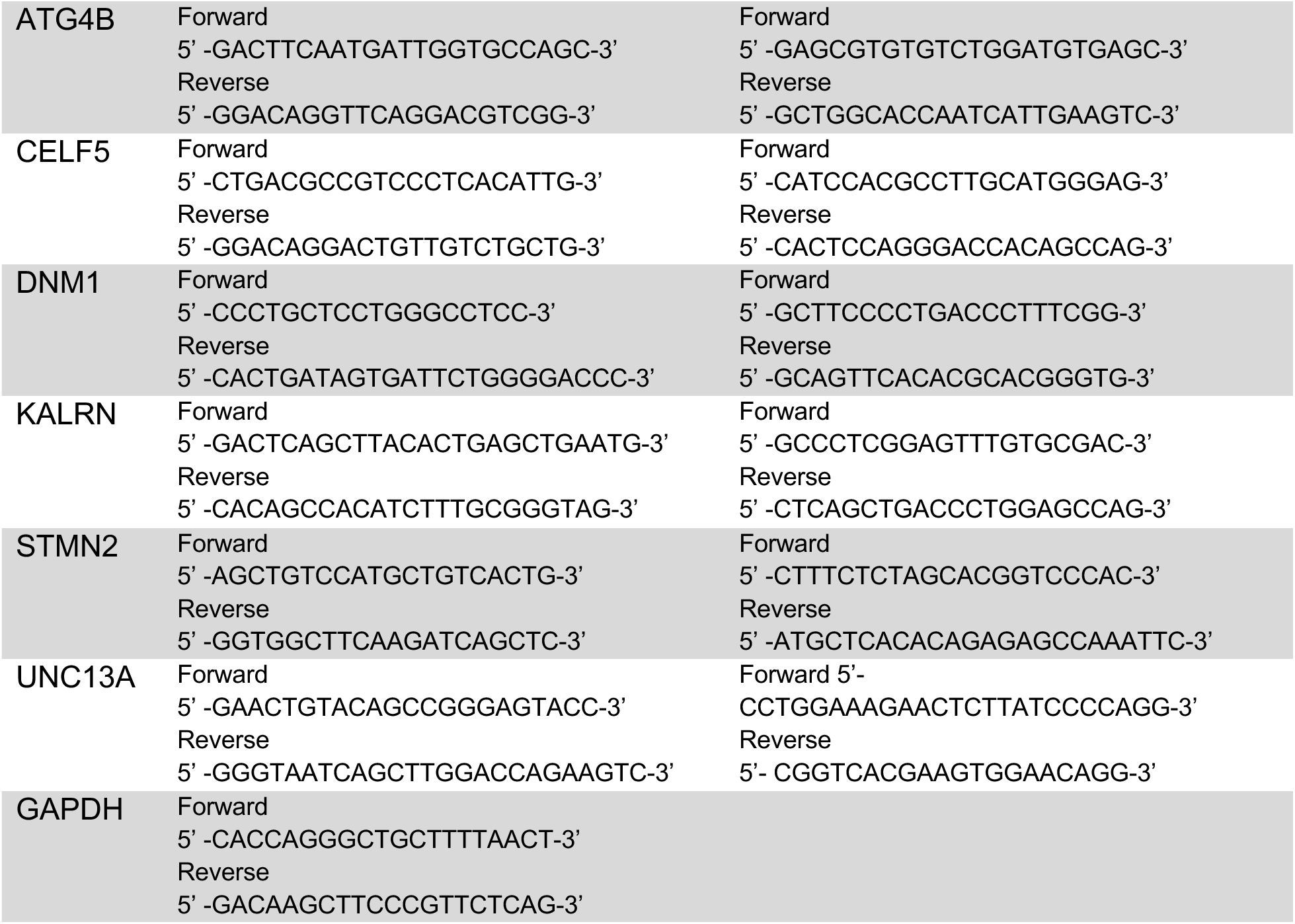
RT-qPCR Primers.

### Western Blot

i^3^Neuron were lysed in RIPA buffer containing 25 mM Tris-HCl pH 7.6, 150 mM NaCl, 1% NP-40, 1% sodium deoxycholate, and 0.1% SDS (Thermo Fisher, PI89900) with protease inhibitor tablet (Sigma-Aldrich, 5892970001). Cell lysates were sonicated, and equal amounts of protein (as measured by DCA assay) were heated at 95°C for 5 minutes in Laemmli buffer with 335 mM BME (Bio-Rad). Protein samples were resolved by SDS-PAGE on 4% - 20% precast gel (Bio-Rad, 4561096) and transferred to 0.2 μm nitrocellulose membrane (Bio-Rad). Membrane were blocked with blocking buffer (Bio-Rad, 12010020) and probed with primary antibodies Anti-GAPDH (Sigma-Aldrich, G9545; 1:5000) (Thermo Fisher, AM4300: 1:5000), Anti-RPL24 (Proteintech, 17082-1-AP; 1:1000), Anti-ABLIM1 (Proteintech, 15129-1-AP; 1:1000), Anti-ITM2C (Abcam, ab101389; 1:1000), Anti-STMN1 (Cell Signaling, 3352T; 1:1000), Anti-BAIAP2 (Proteintech, 11087-2-AP; 1:600), Anti-GABARAP (Proteintech, 18723-1-AP; 1:200), Anti-GABARAPL2 (Abcam, ab126607; 1:400), and anti-TDP-43 (Abcam AB104223, clone 3H8; 1:5000) (Proteintech, 10782-2-AP; 1:1000) for overnight or up to 48 hours of incubation at 4°C. Blots were washed and probed with corresponding secondary antibodies IRDye 800 Donkey anti-rabbit (Licor, 926-32213; 1:20000) and IRDye 680 Donkey anti-mouse (Licor, 926-68022; 1:20000) at room temperature for 1 hour. Blots were visualized on LI-COR Odyssey CLx Imager.

### Western Blot Quantification

Fluorescent signal quantification from the blots was performed using LI-COR Image Studio software. For each lane, rectangles of identical size were drawn around bands of the expected molecular weight to measure signal intensity. For each blot, nearby background signal was quantified for subtraction from signal intensity. For each lane, signal was normalized to protein intensity of an internal loading control (either GAPDH or RPL24). Normalized values were plotted using Graphpad Prism. For purposes of measuring STMN1 levels, we only report on experiments where the average STMN2 levels (in either sgTDP-43 or sgSTMN2 samples) showed at least a 96% reduction.

### Immunofluorescence

For Immunofluorescence, 50000 i^3^Neuron were plated per well onto a PLO-coated glass-bottom 96-well plate (Revvity, 6055302) at Day 3 and maintained as described above. At Day 17, cells were fixed with 4% paraformaldehyde in PBS for 10 minutes at room temperature. Following fixation, cells were washed three times with PBS and permeabilized with 0.5% Triton X-100 (Sigma-Aldrich, X-100) in PBS for 10 minutes at room temperature. Cells were then blocked in 3% BSA + 0.05% Triton X-100 in PBS for 1 hour at room temperature followed by primary antibody incubation at 4 °C overnight. The primary antibodies and dilution used were: Anti-TDP-43 (Proteintech, 10782-2-AP; 1:800) and Anti-TUBB3 (Biolegend, 801201; 1:500). After probing with primary antibody, cells were incubated with secondary antibodies at room temperature for 1 hour as follows: CF488A Donkey Anti-Rabbit IgG (Biotium, 20015; 1:4000) and CF640R Goat Anti-Mouse IgG2a (Biotium, 20261; 1:4000). Images were acquired using Nikon CSU-W1 Spinning Disk confocal microscope with 60X water objective (Nikon).

### RNA sequencing, differential gene expression, and differential splicing analysis

RNA library preparation and sequencing of i^3^Neurons were performed as previously described, and differential gene expression results (using DESeq2^83^ as previously described) were acquired from Brown et al. source data^11^. Unlike in our previous study, we elected to use a Benjamini-Hochberg (BH)-adjusted p-value < 0.05 threshold to designate genes as differentially expressed. Differential splicing analysis was performed using a combination of two computational tools: MAJIQ^84^ and rMATS turbo^85^. As previously described, MAJIQ (v. 2.1, as implemented in a custom Snakemake pipeline^11^) was used to identify junctions with differential splicing and to determine percent spliced in (PSI, Ψ) values and corresponding probabilities of change. To detect additional alternative cassette exon splicing events, we ran rMATS (v. 4.1.2) on STAR aligned BAMs, using the following settings: (--gtf Homo_sapiens.GRCh38.103.gtf -t paired --readLength 75 --nthread 30 --novelSS --mel 1500).

Significant differential splicing events were defined as those meeting the following criteria: for MAJIQ, absolute deltaPSI (ΔΨ) > 0.1 and probability(ΔΨ) > 0.9; for rMATS skipped exon results, absolute IncLevelDifference > 0.1 and false discovery rate (FDR) < 0.1. To identify “cryptic” splicing events, we adapted criteria used in previous studies, but more broadly defined such junctions/events as having significantly increased usage relative to low baseline levels in control samples. This criteria allowed us to identify additional events that may be cryptic in the context of i^3^Neurons, regardless of their splicing patterns in other cell types or contexts (similarly to ref.^12^). Specifically, cryptic splicing events were defined using the following criteria: for MAJIQ, absolute ΔΨ > 0.1, probability(ΔΨ) > 0.9, and control PSI < 0.05; for rMATS, absolute IncLevelDifference > 0.1, FDR < 0.1, and IncLevel_control < 0.05 or > 0.95.

### Proteomics sample preparation, liquid chromatography and mass spectrometry (i^3^Neurons)

i^3^Neurons were grown on 6-well plates (2 wells per replicate; sgControl N=6 and sgTDP-43 N=5) and harvested on day 17 post doxycycline-induction of NGN2. Cells were first washed with cold PBS, and then proteins were extracted using high-percentage detergent lysis buffer on ice, as previously reported.^86^ In short, 200ul of extraction buffer was added to each well, containing 50mM HEPES, 50mM NaCl, 5mM EDTA, 1% SDS, 1% Triton X-100, 1% NP-40, 1% Tween 20, 1% deoxycholate, 1% glycerol, supplemented with cOmplete protease inhibitor tablets (1 tablet per 10mL buffer). Then 10mM dithiotreitol was added to reduce the lysate on a ThermoMixer with 1200 rpm shaking at 60C for 30 min, followed by adding 20mM iodocaetamide alkylation at room temperature in dark for 30 min. Hydrophilic magnetic beads were used to capture the denatured proteins for on-bead tryptic digestion on a ThermoMixer with 1200 rpm shaking at 37C for 30 min.

The tryptic peptides were subjected to liquid chromatography and mass spectrometry (LC-MS) analysis. Peptides were injected into an UltiMate 3000 nano-HPLC and analyzed on an Orbitrap Eclipse MS. Briefly, the samples were separated on a ES903 nano column (75 μm × 500 mm, 2 μm C18 particle) using a 2-h efficient linear gradient with constant flow rate of 300 nL/min with 2%–35% phase B containing 5% DMSO in 0.1% formic acid of acetonitrile. Data was acquired using the data independent acquisition (DIA). MS1 resolution was set to 120K scan range from 400-1000 m/z. For DIA scans, the isolation window was set to 8 m/z and MS2 resolution was 30K and 30% high collision energy. The loop control was set to 3 seconds to maximize the scan cycle.

### Proteomics database search and quality control analysis

The MS raw files were generated (6 control samples and 5 TDP-43 knockdown samples) and database searched against Uniprot-Human-Proteome_UP000005640 using the direct-DIA mode in Spectronaut (v14.1). The false discovery rate of peptide and protein were set to 1%, and mis-cleavage was set to 2, and no medium normalization was selected. For data quality control analysis, the following criteria were used for protein/peptides: a) FDR<0.01; b) Unique peptide >=1; c) Cannot be a common MS contamination; d) cannot be matched to reverse database(decoy); f) have to be identified in more than half of the samples (i.e., # of identification>6 out of 12); g) missing values were assigned as blank (not zero intensity). All missing values were kept blank. The workflow for results generation was as follows: the raw intensity of each protein was normalized against the total protein intensity identified in the same sample.

### Differential and enrichment analyses of i^3^Neuron proteomics data

Normalized intensity values were further processed by log2 normalization and variance stabilizing transformation, as implemented in the DEP package (v.1.22.0)^87^. Proteins with high missingness (using thr=0 as implemented in filter_missval() and groups defined based on KD conditions) were removed before downstream analyses. No imputation was performed. Differential analysis of KD versus control samples was performed with limma (v.3.56.2)^88^, and differentially expressed proteins (DEPs) were defined as those with BH-adjusted p-values < 0.05. Enrichment of GO terms^25^ was performed using clusterProfiler (v.4.8.2)^89^ and msigdbr (v.7.5.1)^24^. In Figure 1 and Figure S3, filtering was applied to exclude redundant enriched terms (those with high Jaccard similarity to more significantly enriched terms) before visualization, using the following thresholds: Fig. 1E: 0.5; Fig. S2C: 0.3.

### Affinity purification mass spectrometry

iPSCs containing an endogenously GFP tagged TDP-43 (N=8) or negative control iPSCs without the tag (N=8) were differentiated into neurons and harvested on day 17 post dox-induction. A GFP nanobody pull-down was performed both by manual operation (N=4 for GFP-tagged and parental line control), and by KingFisher robotic protocol (N=4 for GFP-tagged and parental line control). On-bead tryptic digestion was then performed and 1ug of peptides were subjected to LC-MS analysis and run in data dependent acquisition (DDA) mode where the top 20 most abundant MS1 peaks per acquisition cycle were subjected to MS2 analysis. The database search of RAW files was conducted using Proteome Discoverer (v2.5).

Resulting protein intensities were filtered, log2-normalized and transformed using DEP as described earlier for the manual and KingFisher generated datasets individually to account for preparation-related batch effects. Differential analysis was performed with limma to generate nominal and BH-adjusted p-values and log2 fold change (LFC) values. Proteins detected in at least one GFP-tagged sample and no control samples were assigned positive LFC values and adj. p-value = 1e-2. Significant interactors were defined as those with adjusted p-values < 0.05 and LFC > 0, for either preparation method. For downstream integration and network analyses, these interactors were combined with other experimentally determined TDP-43 interactors as described in “Protein-protein/protein-metabolite interactome construction” (Table S2).

### Proteomics sample preparation, liquid chromatography and mass spectrometry (C9-FTLD patient samples)

Postmortem brain samples from C9orf72-FTLD-TDP patients and cognitively healthy individuals were obtained from the Mayo Clinic Florida Brain Bank as previously described^66^. Briefly, for each sample, 50 mg of snap frozen brain was homogenized using 1ml Mammalian Protein Extraction Reagent (Thermo Scientific, Cat # 78501) with cOmplete protease inhibitor on a TissueLyser II (Qiagen) with 10 seconds per cycle for 3 cycles. The supernatants were collected after centrifugation at 4C with 20,000 g for 30 minutes. A standard BCA Protein Assay (Thermo Scientific, Cat # 23225) was performed to determine the protein concentration: 40 µg normalized protein for each sample in 100 µl of the extraction buffer for the whole cell proteome (described above). The following steps are identical to the whole-cell proteome. The resulting tryptic peptides were analyzed on LC-MS using the DIA approach. The database search was carried out using Spectronaut (v14.1) as described above.

### Differential and enrichment analyses of C9-FTLD patient proteomics data

Quantified protein intensity values were pre-processed and normalized using DEP functions as described above for i^3^Neuron proteomics. A more lenient threshold was set for the initial protein filtering step to retain a greater proportion of proteins for downstream analysis (filter_missval(thr = 5)). Differential analysis comparing C9-FTLD and healthy control samples was performed using limma.

### Analysis of published TDP-43 CLIP data

Genomic binding site data (hg38) from TARDBP/TDP-43 CLIP experiments was downloaded from the POSTAR3^17^ CLIPdb module and filtered to include only data from the following neuron-related cell lines/tissues: brain, hippocampus, frontal cortex, cortical, and SH-SY5Y. Binding sites were annotated and assigned to genes with the HOMER (v.4.9.1)^90^ annotatePeaks.pl function.

### Gene set enrichment analysis of protein/RNA discordance

All genes with estimated RNA and protein LFC values (from DESeq2 and limma respectively) were assigned a discordance score defined as:

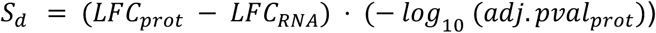

Gene set enrichment analysis (GSEA)^91^ as implemented in clusterProfiler using MSigDB curated GO genesets was performed on proteins ranked by discordance score, after excluding those meeting the following criteria:

1. Protein has differential splicing in i^3^Neurons (i.e. protein in DEP + DS set)
2. Protein is in DEP + DEG, no DS set, and LFCprot and LFCRNA have same sign (protein has concordant RNA expression change)

### Generalized linear model

Absolute LFC values for DEPs were modeled as a function of binary predictor variables indicating the presence of differential gene splicing, differential gene expression, evidence of TDP-43 RNA binding (CLIP), and evidence of TDP-43 protein interaction using a generalized linear model with gamma distribution and log link function. All pairwise interaction terms were also included in the model. Coefficients, standard errors, and corresponding confidence intervals were estimated using the glm function in R (v. 4.3.1).

### Analysis of published splicing analyses

Previously published alternative splicing and polyadenylation datasets were downloaded as described in Table S5. Differentially spliced (“mis-spliced”) and “cryptic” splicing events and genes were determined using study-specific criteria detailed in Table S5; for most analyses, these criteria were defined in the original studies. Certain studies (such as Ma et al. 2022) reported only high-confidence lists of genes, which were described as having alternative splicing and/or cryptic splicing/polyadenylation; these lists were not filtered by any additional criteria. As noted earlier, analyses can differ substantially in their criteria for “cryptic” splicing events, but nearly all studies define cryptic splicing as a subset of all mis-splicing events. As such, for studies that reported only cryptic splicing events, all corresponding genes were counted as having both “mis-splicing” and “cryptic splicing” in reported dataset counts (Fig. S4). Some genes include multiple well-documented junctions with differential or cryptic splicing; for the purposes of this gene-level compendium, we did not differentiate between genes with one vs. multiple mis-splicing events of the same category.

### Inference of differential transcription factor (TF) activity

Differential transcription factor (TF) activity was inferred using a univariate linear model (ULM) applied to t-statistics from the differential gene expression analysis (TDP-43 KD vs. control), as implemented in decoupleR (v.1.8.0)^92^ with regulon sets for 1186 TFs provided by CollecTRI^38^. Genes showing evidence of significant, high magnitude differential splicing (|IncLevelDifference| > 0.25 or |ΔΨ| > 0.25) were excluded prior to analysis (with 15,488 genes remaining). Resulting p-values were corrected for multiple testing using the Benjamini-Hochberg (BH) procedure. TFs with BH-adjusted p-values < 0.1 were designated as significantly differentially active (DA). Regulon sets for DA TFs were intersected with DEGs and tested for enrichment of GO terms with clusterProfiler. For visualization in Fig. S5B, redundant enriched terms were filtered out as described earlier (“Differential and enrichment analyses of proteomics data”), using a threshold of 0.25.

### RNA binding protein motif enrichment analysis

RBP motif enrichment was assessed using rMAPS2^45^ on rMATS-identified skipped exon events (hereafter referred to as misspliced exons, MEs). Analyses were performed on three sets of ME events (Fig S6A):

1. **DS Set 1**: all ME events
2. **DS Set 2**: ME events in which the region between the upstream exon end and downstream exon start showed no overlap with reported TDP-43 binding sites as defined in “Analysis of published TDP-43 CLIP data”. These ME events were further filtered to exclude those overlapping TDP-43 binding sites reported in a secondary iCLIP dataset^18^. (Processed iCLIP peaks from this study were downloaded from the iCount server; http://icount.biolab.si/, download date: May 11, 2023.)
3. **DS Set 3**: ME events from DS Set 2 in which none of a subset of canonical TDP-43 motifs (defined below) was detected in the regions surrounding the cassette exon, using rMAPS default definitions of upstream and downstream windows (250 bp upstream and downstream, including 50 bp into exon coordinates: −50 to +250 relative to the upstream exon end; −250 to +50 relative to the downstream exon start).

The rationale for this design was as follows: DS Set 1 captures motifs enriched around all misspliced exon events, including those potentially regulated by TDP-43. This allows detection of RBPs that may act cooperatively or competitively with TDP-43. DS Sets 2 and 3 exclude splice events with reported and/or predicted evidence of nearby TDP-43 binding, thereby allowing us to detect motifs and RBPs more likely to act independently of TDP-43. Genomic interval operations described above were performed with PyRanges (v.0.0.127).

Previously described TDP-43 binding motifs^36,46^ used in the analysis are listed in Table 3 below:

**Table 3:**
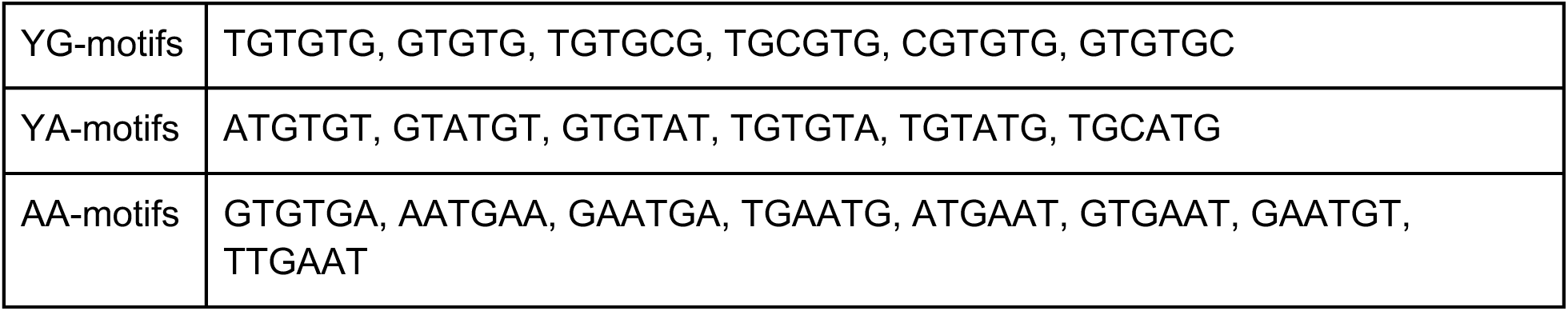
TDP-43 binding motif groups.

The subset of motifs selected for exclusion in DS Set 3 are TGTGTG, GTGTGT, TGCGTG, CGTGTG, ATGTGT, TGTGTA, TGCATG, GTGTGA, based on significance of motif enrichment patterns detected in DS Sets 1/2 (Fig. S6C). Additional RBP motifs for rMAPS2 analysis were obtained from the ATtRACT database^93^ and preprocessed and simplified with a custom R script.

Significance of enrichment was determined by retaining, for each motif-RBP pair, the minimum p-value across all intronic/exonic regions tested in rMAPS2 (upstream exon 3’ end, upstream intron 5’ end, upstream intron 3’end, ME 5’ end, ME 3’ end, downstream intron 5’ end, downstream intron 3’ end, downstream exon 5’ end). These values were Bonferroni-corrected, and significant motif enrichment results were defined based on the following adjusted p-value thresholds (DS Set 1: 1e-4, DS Set 2: 1e-3, DS Set 3: 1e-3). Motif-RBP pairs were further manually curated and filtered based on plotted motif enrichment maps, as generated by rMAPS2. Additional motif enrichment maps with nominally significant p-values were also visualized for RBPs associated with secondary motifs meeting stricter enrichment criteria.

### Protein-protein/protein-metabolite interactome construction

The reference interactome of protein-protein and protein-metabolite interactions for use in OmicsIntegrator2 was generated using a combination of publicly available datasets and significant interactors inferred from AP-MS experiments in this study (“Affinity purification mass spectrometry”). Briefly, experimentally validated protein-protein interactions were downloaded from iRefIndex^94^ v.19, IntAct^95^, BioGRID^96^, and supplemented with brain-specific interactions determined by BraInMap^97^, defining edge costs as previously described^53,55^. Protein-metabolite interactions were derived from HMDB and Recon 2, also as previously described^56^.

### Network optimization and analysis

Network optimization was performed using the Prize-Collecting Steiner Forest (PCSF) algorithm as implemented in OmicsIntegrator2 (v.2.4.0.1) (https://github.com/fraenkel-lab/OmicsIntegrator2/tree/dev-py3.9). This method aims to generate a high-confidence subnetwork of protein-protein interactions that prioritizes proteins (nodes) with experimental significance or literature relevance to a specific context. Briefly, the approach projects a set of input nodes with user-assigned prize values onto a reference interactome and identifies an optimized, connected subnetwork (“solution network”) that maximizes the weighted sum of retained prizes while minimizing the total cost of included edges. Three hyperparameters govern the optimization: beta (b), which scales the prizes and influences overall network size; omega (w), which controls the number of individual trees included in the solution; and gamma (g), which penalizes highly connected hub nodes. Complete mathematical details, including the formalized PCSF optimization objective, are provided in Tuncbag et al. (2016)^52^ and Phatnani et al. (2021)^53^.

For our analysis, input nodes included differentially expressed proteins, inferred TFs, inferred RBPs, and differentially spliced genes that were differentially expressed at the RNA level but not sufficiently detected in proteomics. Node prizes were assigned as follows: DEPs: absolute log2 fold change; inferred TFs: −log10 adjusted p-value for differential activity; inferred RBPs: −log10 adjusted p-value for motif enrichment (individually computed for each DS set); DEGs with differential splicing: absolute log2 fold change in gene expression. Within each analysis, all prize values were normalized to the 0-1 range prior to compilation. For genes assigned multiple prize values (i.e. reached significance across multiple analyses), the highest normalized value was kept.

To assess robustness and specificity of network solutions, we applied the following randomization procedure. First, Gaussian noise was added to interactome edge weights across 100 randomizations, and node robustness was calculated as the fraction of randomizations in which a node was included in the solution. Second, node prizes were randomly reassigned to interactome nodes across 100 randomizations, and inverse specificity was defined as the fraction of randomizations in which a node was recovered. “Robust-filtered” networks were generated by retaining only nodes with robustness > 0.6 and inverse specificity < 0.4.

Hyperparameter tuning was performed by testing multiple parameter sets (b = {1, 5, 10, 20}, w = {1, 3, 5, 10}, g = {5, 6, 7}) and selecting the combination that resulted in a combination of minimal mean network inverse specificity, maximal mean robustness, minimal Kolmogorov– Smirnov statistic between the degree distributions of prize nodes and solution nodes, and that also retained at least 60% of input nodes, for robust-filtered solution networks. The final parameters were set to b = 10, w = 10, g = 5.

The corresponding robust-filtered solution network consisted of 723 nodes. Leiden community detection (resolution = 1) was performed with Python package igraph (v.0.11.4) to identify network clusters. GO term enrichment analyses were performed both on the full network and on individual clusters using clusterProfiler and MSigDB curated GO genesets. The clustered and annotated network was visualized in Cytoscape (v.3.9.1)^98^.

### Differential analysis of previously published post-mortem datasets

Published post-mortem proteomic and transcriptomic datasets, with corresponding sample metadata, were downloaded as detailed in Table S14. Each dataset was processed separately using DEP and limma (proteomics) or DESeq2 (RNA-seq), with inclusion of relevant study-specific covariates where applicable.

For the single-cell proteomics dataset of Guise et al. (2024)^68^, which profiled motor neurons stratified by TDP-43 pathology status, differential analysis was performed across four contrasts: ALS neurons with no, mild, moderate, or severe pathology versus control neurons. Donor ID was used as a covariate for all differential analyses, following recommendations outlined in the original study. For the NYGC RNA-seq dataset, analyses were stratified by tissue region and disease subtype annotations defined previously^11^. To minimize technical confounding, only Novaseq-sequenced samples were included, retaining a total of 745 samples. The following contrasts were performed, including sex and processing site as covariates:

● ALS-TDP vs. control: frontal cortex, motor cortex, cervical spinal cord, and lumbar spinal cord.
● FTLD-TDP vs. control: frontal cortex and temporal cortex.

GO term enrichment analyses were performed on sets of differentially expressed genes and proteins using msigdbr and clusterProfiler. Enrichment results were visualized in dotplot format; for pathways enriched among both upregulated and downregulated genes or proteins, the more significant enrichment was reported.

Heatmaps of gene LFC values were generated using ComplexHeatmap (v.2.16.0)^99^, with default hierarchical clustering parameters where applicable. In Fig. 5B, genes were included if they showed significant differential expression in at least 2 of both the post-mortem RNAseq and proteomics comparisons, or at least 4 of the post-mortem RNAseq comparisons, or at least 4 of the post-mortem proteomics comparisons. Additional genes of interest (e.g. STMN2) not meeting this criteria were also visualized, as indicated in Fig. 5 legend.

## Data and code availability

Processed RNA-seq and proteomics data is provided as source data on Github repository (see below). Data from previously published studies is available from sources described in Methods and Tables S5 and S14. RNAseq data was downloaded from Gene Expression Omnibus: GSE126543 (Liu et al., 2019), GSE137810 (New York Genome Center). Proteomics data was downloaded from https://www.synapse.org/Synapse:syn10143061 (Umoh et al., 2018) and https://doi.org/10.1016/j.celrep.2023.113636 (Guise et al., 2024). Raw mass spectrometry proteomics data from this study have been deposited to the ProteomeXchange Consortium via the PRIDE^100^ partner repository with the dataset identifier PXD068980. The C9-FTLD patient samples were also previously deposited^65^ to the ProteomeXchange Consortium via PRIDE (ProteomeXchange: PXD047657).

An interactive version of the protein interactome subnetwork is available at: https://fraenkel-lab.github.io/tdp43-multiomic-network/network

Analysis code will be available on Github at https://github.com/fraenkel-lab/tdp43-multiomic-network

## Author contributions

Conceived and designed the experiments: V.K., Z.L., K.B., Y.A.Q., S.R., M.E.W., E.F., S.E.K.-H.

Performed experiments: V.K., Z.L., K.B., Y.A.Q., S.R., M.A., S.E.K.-H.

Analyzed the data: VV.K., Z.L., K.B., Y.A.Q., S.R., S.S., M.A., S.T., A-L.B., P.F., M.E.W., E.F., S.E.K.-H.

Contributed reagents and materials: M.P., L.P, D.W.D., H.J.K, J.P.T. M.E.W., S.E.K.-H.

Wrote the manuscript: V.K., S.E.K.-H.

Edited the manuscript: V.K., Z.L., K.B., Y.A.Q., E.F., S.E.K.-H.

All authors reviewed and approved the final version of this manuscript.

## Conflicts of interest

The authors declare that they have no conflicts of interest.

## Acknowledgments

We thank Steve Coon, James Iben, and Tianwei Li at the NIH/NICHD Molecular Genomics core for RNA sequencing and initial analyses. We thank the NIH/NINDS proteomics core for mass spec facilities. Postmortem human brain tissue was obtained from the Mayo Clinic Florida Brain Bank. We thank the patients and their family for their contributions to research. We thank all members of the Fraenkel lab and Kargbo-Hill lab for helpful discussions. Some Figure subpanels were created in Biorender.

This work was supported by NIH: K99/R00AG080036 (S.E.K.-H.), RF1 AG075901 (E. F.), RF1 AG076214 (E. F.), R01NS120992 (M.P.), U54NS123743 (L.P. and M.P.) This research was funded, in part, by the Intramural Research Program of the National Institutes of Neurological Disorders and Stroke at the National Institutes of Health, USA, (M.E.W) and from the Center for Alzheimer’s and Related Dementias (CARD) at the National Institute on Aging, USA, (Y.Q.) project number ZIAAG000534. The contributions of the NIH authors are considered Works of the United States Government. The findings and conclusions presented in this paper are those of the authors and do not necessarily reflect the views of the NIH or the U.S. Department of Health and Human Services. This work was also funded in part from the Chan Zuckerberg Initiative (M.E.W.). This material is based upon work supported by the National Science Foundation (NSF) Graduate Research Fellowship Program under Grant No. 1745302 (to V.K.).

## Notes

https://www.ndexbio.org/viewer/networks/b343879d-8800-11f0-a218-005056ae3c32

https://github.com/fraenkel-lab/tdp43-multiomic-network

### Competing Interest Statement

The authors have declared no competing interest.

### Summary of Updates

The manuscript has been revision to fix a typo in the link to our integrative network, and other minor textual and figure edits.

