## Supplemental Figures for "Integrative multiomic analysis links TDP-43-driven splicing defects to cascading proteomic disruption of ALS/FTD pathways"

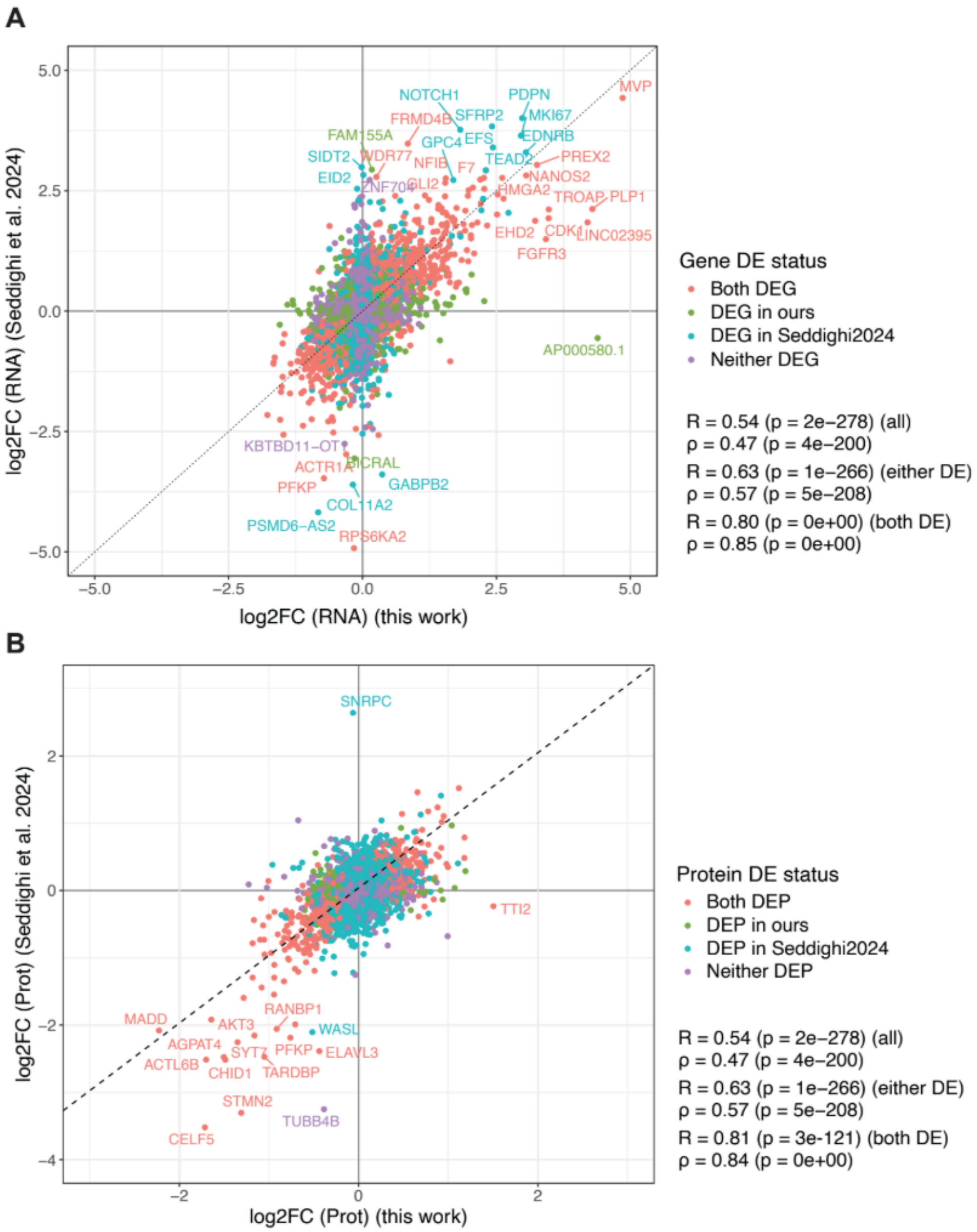


**Figure S1: Correlation between the RNA-seq and proteomics experiments used in this study with prior studies of TDP-43 KD in iNeurons.**
**A)** RNA expression LFC from the iNeuron RNA-seq experiment in Seddighi et al., 2024 plotted against the RNA expression LFC used in this study, which was previously published (Brown et al., 2022). Colors indicate the status of each gene as differentially expressed (DEG) in each study. **B)** Protein expression LFC from Seddighi et al., 2024 plotted against the new proteomics LFC generated in this study. Colors indicate the status of each gene as differentially expressed (DEP) in each study. Pearson (R) and Spearman (rho) correlation values and corresponding p-values for all genes and proteins and differentially expressed subsets indicated.

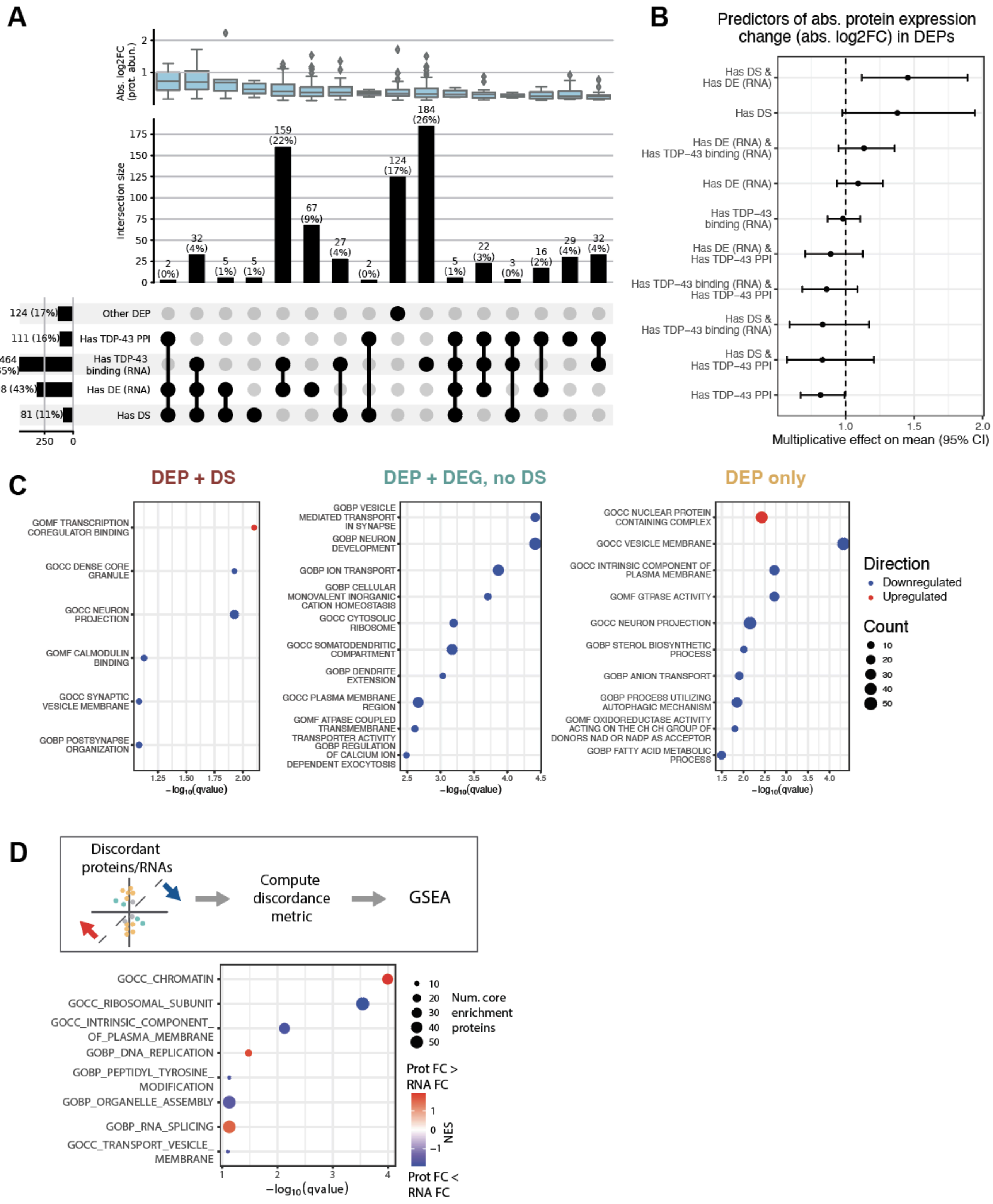
A
Abs. log2FC (prot. abun.)
Intersection size
159 (22%)
184 (26%)
124 (17%)
67 (9%)
32 (4%)
27 (4%)
22 (3%)
29 (4%)
16 (2%)
2 (0%)
5 (1%)
3 (0%)
124 (17%)
111 (16%)
464 (65%)
308 (43%)
81 (11%)
Other DEP
Has TDP-43 PPI
Has TDP-43 binding (RNA)
Has DE (RNA)
Has DS
B
Predictors of abs. protein expression change (abs. log2FC) in DEPs
Has DS & Has DE (RNA)
Has DS
Has DE (RNA) & Has TDP−43 binding (RNA)
Has DE (RNA)
Has TDP−43 binding (RNA)
Has DE (RNA) & Has TDP−43 PPI
Has TDP−43 binding (RNA) & Has TDP−43 PPI
Has DS & Has TDP−43 binding (RNA)
Has DS & Has TDP−43 PPI
Has TDP−43 PPI
Multiplicative effect on mean (95% CI)
C
DEP + DS
GOMF TRANSCRIPTION COREGULATOR BINDING
GOCC DENSE CORE GRANULE
GOCC NEURON PROJECTION
GOMF CALMODULIN BINDING
GOCC SYNAPTIC VESICLE MEMBRANE
GOBP POSTSYNAPSE ORGANIZATION
DEP + DEG, no DS
GOBP VESICLE MEDIATED TRANSPORT IN SYNAPSE
GOBP NEURON DEVELOPMENT
GOBP ION TRANSPORT
GOBP CELLULAR MONOVALENT INORGANIC CATION HOMEOSTASIS
GOCC CYTOSOLIC RIBOSOME
GOCC SOMATODENDRITIC COMPARTMENT
GOBP DENDRITE EXTENSION
GOCC PLASMA MEMBRANE REGION
GOMF ATPASE COUPLED TRANSMEMBRANE TRANSPORTER ACTIVITY
GOBP REGULATION OF CALCIUM ION DEPENDENT EXOCYTOSIS
DEP only
GOCC NUCLEAR PROTEIN CONTAINING COMPLEX
GOCC VESICLE MEMBRANE
GOCC INTRINSIC COMPONENT OF PLASMA MEMBRANE
GOMF GTPASE ACTIVITY
GOCC NEURON PROJECTION
GOBP STEROL BIOSYNTHETIC PROCESS
GOBP ANION TRANSPORT
GOBP PROCESS UTILIZING AUTOPHAGIC MECHANISM
GOMF OXIDOREDUCTASE ACTIVITY ACTING ON THE CH CH GROUP OF DONORS NAD OR NADP AS ACCEPTOR
GOBP FATTY ACID METABOLIC PROCESS
−log10(qvalue)
Direction
Downregulated
Upregulated
Count
D
Discordant proteins/RNAs
Compute discordance metric
GSEA
GOCC_CHROMATIN
GOCC_RIBOSOMAL_SUBUNIT
GOCC_INTRINSIC_COMPONENT_OF_PLASMA_MEMBRANE
GOBP_DNA_REPLICATION
GOBP_PEPTIDYL_TYROSINE_MODIFICATION
GOBP_ORGANELLE_ASSEMBLY
GOBP_RNA_SPLICING
GOCC_TRANSPORT_VESICLE_MEMBRANE
−log10(qvalue)
Num. core enrichment proteins
Prot FC > RNA FC
NES
Prot FC < RNA FC

**Figure S2: Characteristics of DEPs and enriched pathways in TDP-43 KD i$^3$Neurons.**
**A)** Upset plot indicating numbers of DEPs with differential splicing, differential gene expression, evidence of TDP-43 RNA binding and evidence of protein-protein interaction with TDP-43. The upper boxplot shows distributions of absolute LFC protein expression values for intersection sets. **B)** Coefficients and confidence intervals derived from generalized linear model of abs. LFC protein expression values with select gene characteristics of DEPs (as in A) as predictors (see Methods). **C)** Top significantly enriched MSigDB GO terms for protein sets as defined in Fig 1B,C. **D)** (Top) Schematic of gene set enrichment analysis framework for proteins without DS or concordant RNA expression changes. (Bottom) Pathways with significant enrichment among genes with discordant RNA/protein changes.

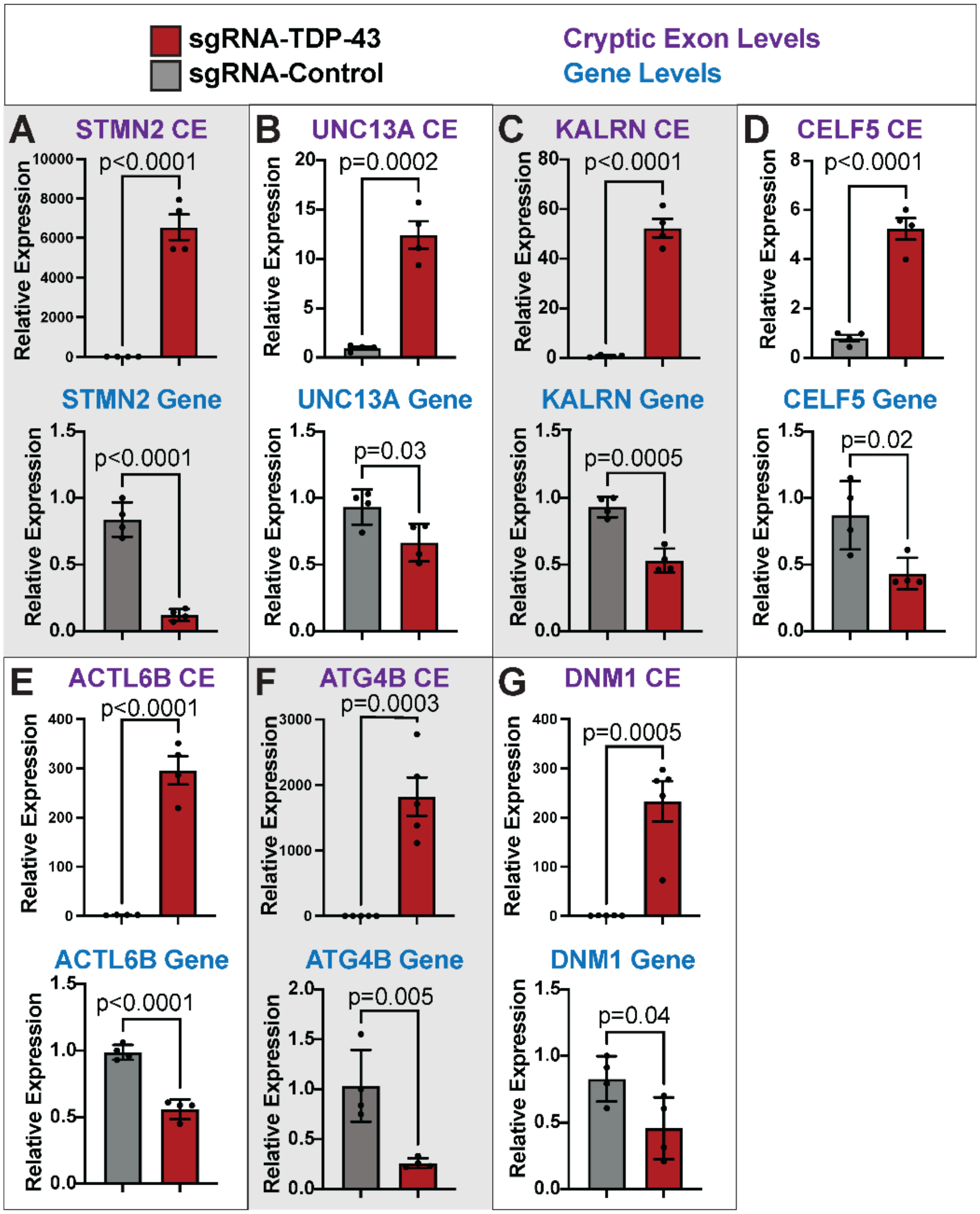


**Figure S3: Validation of cryptic exon expression and decreased transcript levels in TDP-43 KD i3Neurons.**

**A-G)** Quantification of relative expression levels of cryptic exon expression (top) and gene expression (bottom) in Control and TDP-43 KD $i^3$Neurons for a subset of differentially expressed genes which are predicted to express cryptic exons by RNA-seq. Statistical comparison between Control and TDP-43 KD using unpaired t test; Error bars show SEM (top, CE) or SD (bottom, gene level).

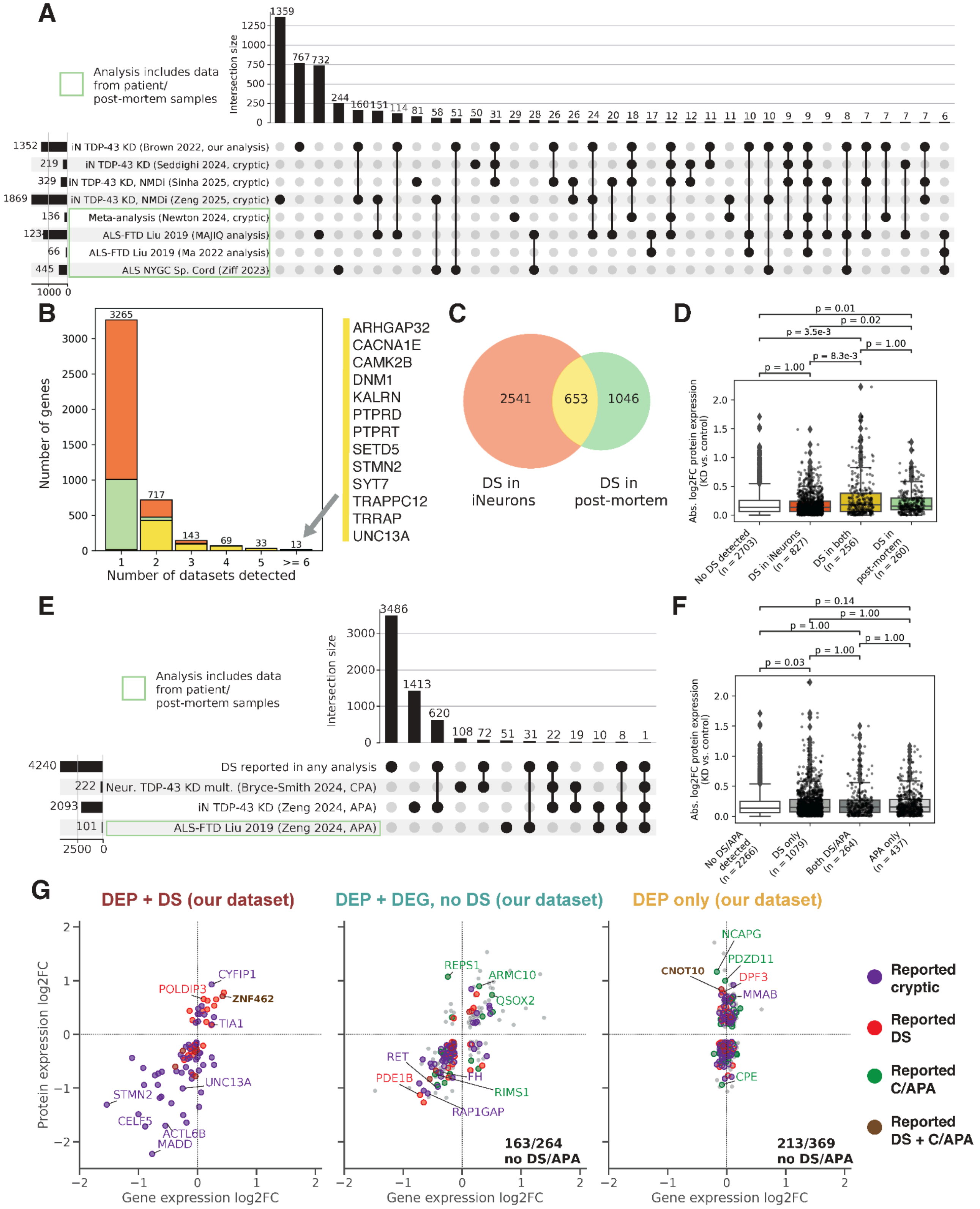
A
Analysis includes data from patient/ post-mortem samples
Intersection size
1359
767
732
244
160
151
114
81
58
51
50
31
29
28
26
26
24
20
18
17
12
12
11
11
10
10
9
9
9
8
7
7
7
7
6
1352
iN TDP-43 KD (Brown 2022, our analysis)
219
iN TDP-43 KD (Seddighi 2024, cryptic)
329
iN TDP-43 KD, NMDi (Sinha 2025, cryptic)
1869
iN TDP-43 KD, NMDi (Zeng 2025, cryptic)
136
Meta-analysis (Newton 2024, cryptic)
ALS-FTD Liu 2019 (MAJIQ analysis)
66
ALS-FTD Liu 2019 (Ma 2022 analysis)
445
ALS NYGC Sp. Cord (Ziff 2023)
1000 0
B
Number of genes
Number of datasets detected
3265
717
143
69
33
13
ARHGAP32
CACNA1E
CAMK2B
DNM1
KALRN
PTPRD
PTPRT
SETD5
STMN2
SYT7
TRAPPC12
TRRAP
UNC13A
C
2541
653
1046
DS in iNeurons
DS in post-mortem
D
Abs. log2FC protein expression (KD vs. control)
p = 0.01
p = 0.02
p = 3.5e-3
p = 1.00
p = 8.3e-3
p = 1.00
No DS detected (n = 2703)
DS in iNeurons (n = 827)
DS in both (n = 256)
DS in post-mortem (n = 260)
E
Analysis includes data from patient/ post-mortem samples
Intersection size
3486
1413
620
108
72
51
31
22
19
10
8
1
4240
DS reported in any analysis
222
Neur. TDP-43 KD mult. (Bryce-Smith 2024, CPA)
2093
iN TDP-43 KD (Zeng 2024, APA)
101
ALS-FTD Liu 2019 (Zeng 2024, APA)
2500 0
F
Abs. log2FC protein expression (KD vs. control)
p = 0.14
p = 1.00
p = 1.00
p = 1.00
p = 1.00
p = 0.03
No DS/APA detected (n = 2266)
DS only (n = 1079)
Both DS/APA (n = 264)
APA only (n = 437)
G
DEP + DS (our dataset)
DEP + DEG, no DS (our dataset)
DEP only (our dataset)
Protein expression log2FC
Gene expression log2FC
CYFIP1
POLDIP3
ZNF462
TIA1
UNC13A
STMN2
CELF5
ACTL6B
MADD
REPS1
ARMC10
QSOX2
RET
PDE1B
FH
RIMS1
RAP1GAP
163/264 no DS/APA
NCAPG
PDZD11
CNOT10
DPF3
MMAB
CPE
213/369 no DS/APA
Reported cryptic
Reported DS
Reported C/APA
Reported DS + C/APA

**Figure S4: Compendium of differential splicing and alternative polyadenylation associated with TDP-43 proteinopathy from previously published studies.**
**A)** Upset plot of 4240 genes with reported differential splicing (DS) (inclusive of cryptic splicing studies) across 8 analyses, including analysis of i3Neuron data presented in this study (listed first). Intersection sets with < 6 genes not shown. **B)** Bar chart indicating number of datasets in which genes are detected as differentially spliced. Colors indicate proportions detected in i3Neurons only (orange), post-mortem samples only (green), or both (gold). **C)** Venn diagram indicating genes reported DS in i3Neuron vs. post-mortem studies. **D)** Box plot of absolute log2 fold changes (LFC) in protein expression across intersection sets defined as in C. Numbers of genes with quantified protein expression from proteomics are indicated. Statistics indicate Bonferroni-corrected p-values from two-sided Mann-Whitney-Wilcoxon tests. **E)** Upset plot of 5841 genes with reported differential splicing (as in A) or cryptic/alternative polyadenylation across 3 C/APA datasets. **F)** Box plot of absolute LFC in protein expression across genes reported to have either DS or C/APA or both. Numbers of genes with quantified protein expression from proteomics are indicated. Statistics indicate Bonferroni-corrected p-values from two-sided Mann-Whitney-Wilcoxon tests. **G)** Protein expression LFC plotted against RNA expression LFC for DEP groups as defined in Fig 1D. Colors indicate genes reported to have cryptic or differential splicing (DS), or cryptic/alternative polyadenylation (C/APA) across collected datasets. Previously studied and notable genes of interest are labeled.

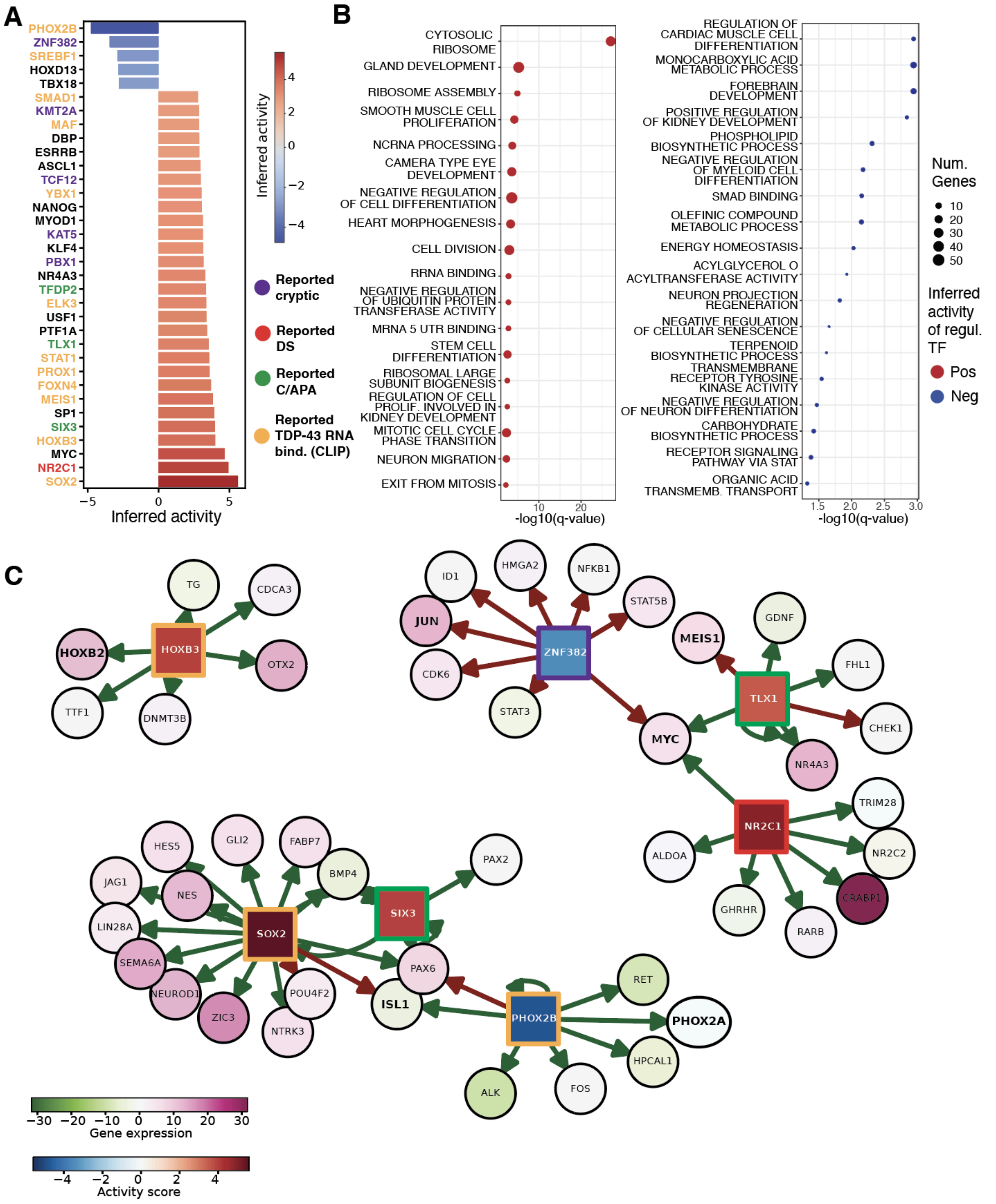

A
PHOX2B
ZNF382
SREBF1
HOXD13
TBX18
SMAD1
KMT2A
MAF
DBP
ESRRB
ASCL1
TCF12
YBX1
NANOG
MYOD1
KAT5
KLF4
PBX1
NR4A3
TFDP2
ELK3
USF1
PTF1A
TLX1
STAT1
PROX1
FOXN4
MEIS1
SP1
SIX3
HOXB3
MYC
NR2C1
SOX2
−5
0
5
Inferred activity
Inferred activity
4
2
0
−2
−4
Reported cryptic
Reported DS
Reported C/APA
Reported TDP-43 RNA bind. (CLIP)
B
CYTOSOLIC RIBOSOME
GLAND DEVELOPMENT
RIBOSOME ASSEMBLY
SMOOTH MUSCLE CELL PROLIFERATION
NCRNA PROCESSING
CAMERA TYPE EYE DEVELOPMENT
NEGATIVE REGULATION OF CELL DIFFERENTIATION
HEART MORPHOGENESIS
CELL DIVISION
RRNA BINDING
NEGATIVE REGULATION OF UBIQUITIN PROTEIN TRANSFERASE ACTIVITY
MRNA 5 UTR BINDING
STEM CELL DIFFERENTIATION
RIBOSOMAL LARGE SUBUNIT BIOGENESIS
REGULATION OF CELL PROLIF. INVOLVED IN KIDNEY DEVELOPMENT
MITOTIC CELL CYCLE PHASE TRANSITION
NEURON MIGRATION
EXIT FROM MITOSIS
10
20
-log10(q-value)
REGULATION OF CARDIAC MUSCLE CELL DIFFERENTIATION
MONOCARBOXYLIC ACID METABOLIC PROCESS
FOREBRAIN DEVELOPMENT
POSITIVE REGULATION OF KIDNEY DEVELOPMENT
PHOSPHOLIPID BIOSYNTHETIC PROCESS
NEGATIVE REGULATION OF MYELOID CELL DIFFERENTIATION
SMAD BINDING
OLEFINIC COMPOUND METABOLIC PROCESS
ENERGY HOMEOSTASIS
ACYLGLYCEROL O ACYLTRANSFERASE ACTIVITY
NEURON PROJECTION REGENERATION
NEGATIVE REGULATION OF CELLULAR SENESCENCE
TERPENOID BIOSYNTHETIC PROCESS
TRANSMEMBRANE RECEPTOR TYROSINE KINASE ACTIVITY
NEGATIVE REGULATION OF NEURON DIFFERENTIATION
CARBOHYDRATE BIOSYNTHETIC PROCESS
RECEPTOR SIGNALING PATHWAY VIA STAT
ORGANIC ACID TRANSMEMB. TRANSPORT
1.5
2.0
2.5
3.0
-log10(q-value)
Num. Genes
10
20
30
40
50
Inferred activity of regul. TF
Pos
Neg
C
TG
CDCA3
HOXB2
HOXB3
OTX2
TTF1
DNMT3B
ID1
HMGA2
NFKB1
STAT5B
JUN
ZNF382
CDK6
STAT3
MEIS1
GDNF
FHL1
TLX1
CHEK1
MYC
NR4A3
TRIM28
NR2C1
ALDOA
NR2C2
CRABP1
GHRHR
RARB
HES5
GLI2
FABP7
JAG1
NES
BMP4
PAX2
SIX3
LIN28A
SOX2
SEMA6A
PAX6
NEUROD1
POU4F2
ISL1
ZIC3
NTRK3
RET
PHOX2B
PHOX2A
HPCAL1
ALK
FOS
−30
−20
−10
0
10
20
30
Gene expression
−4
−2
0
2
4
Activity score

**Figure S5: Inference of differential TF activity in TDP-43 KD neurons.**
**A)** Bar plot of inferred activity levels for 34 TFs inferred to be significantly differentially active (BH adj. p-value < 0.1) (Methods). Gene name colors indicate evidence of reported cryptic splicing, DS, C/APA, or TDP-43 RNA binding. **B)** Top significantly enriched MSigDB GO terms in DEGs putatively regulated by inferred differentially active TFs. Results for TFs inferred to have increased activity (left) and decreased activity (right) in TDP-43 KD vs. control. **C)** Network plot of top inferred TFs and prominent regulon targets with significant gene expression changes. Arrow color indicates direction of regulation (green: activation, red: suppression). Square nodes indicate inferred TFs and are colored according to inferred activity score. Round nodes indicate gene targets and are colored by scaled RNA expression change (TDP-43 KD vs. control). TF node border colors as in A. Target genes encoding other prominent TFs in bold.

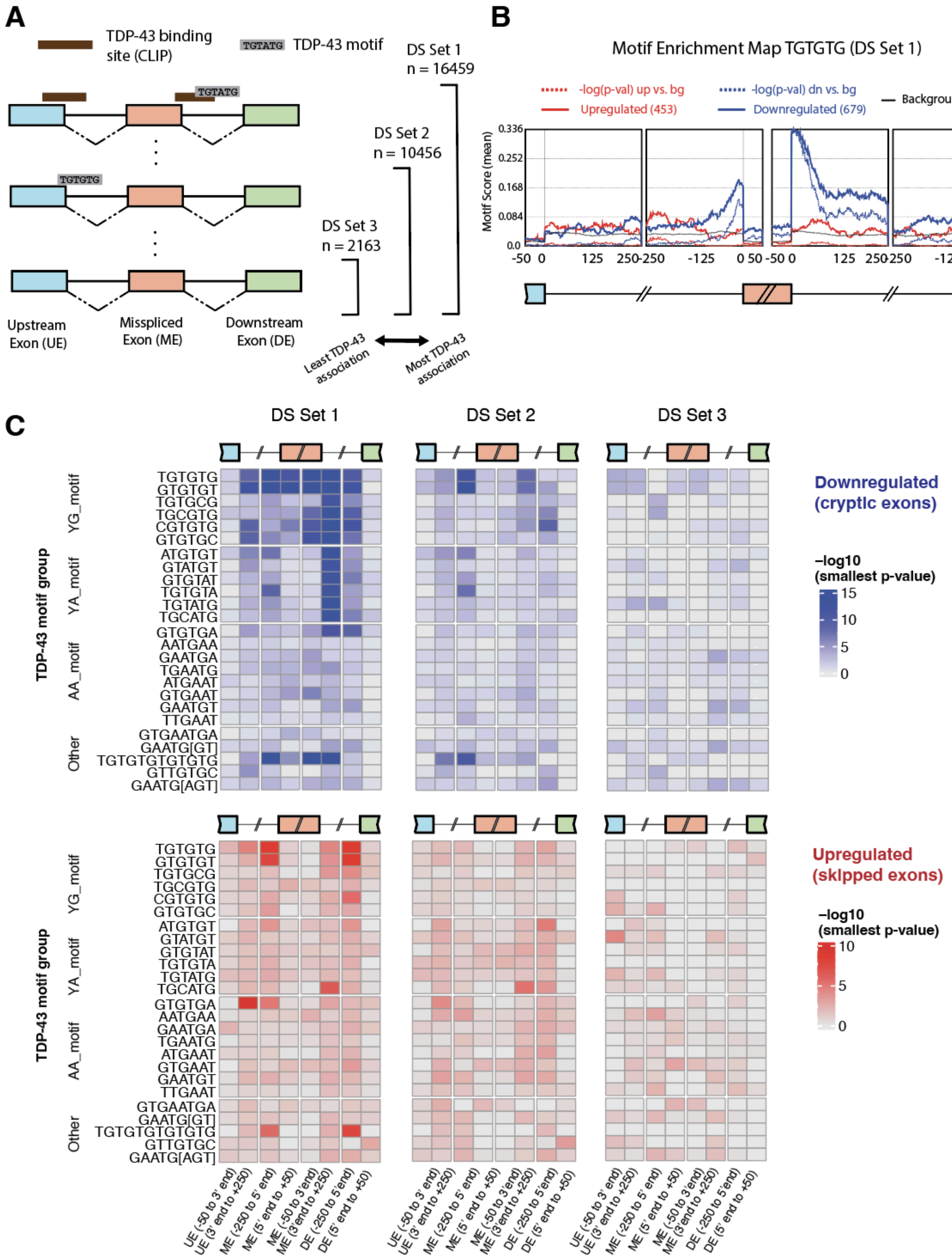
A
TDP-43 binding site (CLIP)
TGTATG TDP-43 motif
DS Set 1 n = 16459
DS Set 2 n = 10456
DS Set 3 n = 2163
Upstream Exon (UE)
Misspliced Exon (ME)
Downstream Exon (DE)
Least TDP-43 association
Most TDP-43 association
B
Motif Enrichment Map TGTGTG (DS Set 1)
-log(p-val) up vs. bg
Upregulated (453)
-log(p-val) dn vs. bg
Downregulated (679)
Background (15327)
Motif Score (mean)
Negative log10(p-value)
C
DS Set 1
DS Set 2
DS Set 3
TDP-43 motif group
YG_motif
YA_motif
AA_motif
Other
Downregulated (cryptic exons)
-log10 (smallest p-value)
Upregulated (skipped exons)
UE (-50 to 3' end)
UE (3' end to +250)
ME (-250 to 5' end)
ME (5' end to +50)
ME (-50 to 3' end)
ME (3' end to +250)
DE (-250 to 5' end)
DE (5' end to +50)

**Figure S6: Analysis of RBPs with enriched binding motifs around misspliced exons.**
**A)** Schematic of criteria used to create different sets of misspliced exon (ME) junctions for RBP motif enrichment testing (Methods). Each successive DS set is generated by excluding junctions near TDP-43 CLIP binding sites (neuronal experiments only) (DS Set 2), and near canonical TDP-43 motifs (DS Set 3). Junction windows tested for overlaps (including upstream and downstream exons and proximal intronic regions) are described in Methods. **B)** Motif enrichment map for TGTGTG motif (prominent motif associated with TDP-43) for DS Set 1 covering the 50 base pairs (bp) at the ends of mis-spliced exon (orange), upstream exon (blue) or downstream exon (green), or in the 250 bp intronic regions proximal to exons. Solid lines indicate motif enrichment score (left y-axis), dotted lines indicate corresponding -log10(p-value) (right y-axis). Motif enrichment is computed separately for exons downregulated (blue) and upregulated (red) in control vs. TDP-43 KD. Evidence of strong motif presence downstream of downregulated (cryptic) exons (and some enrichment immediately upstream) is consistent with previous studies of TDP-43 RNA binding activity. **C)** Heatmaps of minimum motif enrichment p-values across relevant junction regions (Methods) for all canonical TDP-43 motifs (Hallegger et al. 2021) and all DS sets tested.

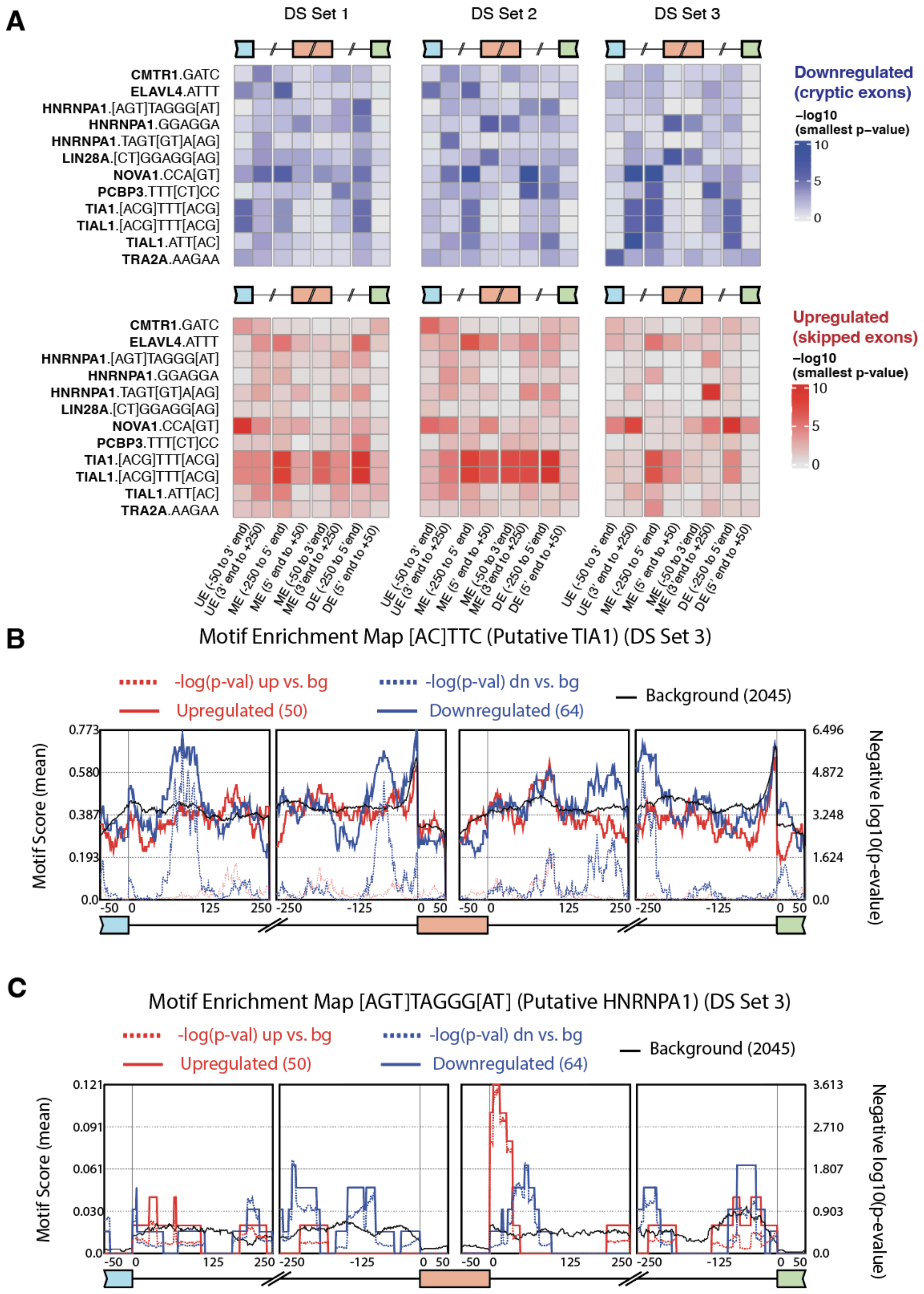
A
DS Set 1
DS Set 2
DS Set 3
CMTR1.GATC
ELAVL4.ATTT
HNRNPA1.[AGT]TAGGG[AT]
HNRNPA1.GGAGGA
HNRNPA1.TAGT[GT]A[AG]
LIN28A.[CT]GGAGG[AG]
NOVA1.CCA[GT]
PCBP3.TTT[CT]CC
TIA1.[ACG]TTT[ACG]
TIAL1.[ACG]TTT[ACG]
TIAL1.ATT[AC]
TRA2A.AAGAA
Downregulated (cryptic exons)
−log10 (smallest p−value)
Upregulated (skipped exons)
UE (-50 to 3' end)
UE (3' end to +250)
ME (-250 to 5' end)
ME (5' end to +50)
ME (-50 to 3' end)
ME (3' end to +250)
DE (-250 to 5' end)
DE (5' end to +50)
B
Motif Enrichment Map [AC]TTC (Putative TIA1) (DS Set 3)
-log(p-val) up vs. bg
-log(p-val) dn vs. bg
Upregulated (50)
Downregulated (64)
Background (2045)
Motif Score (mean)
Negative log10(p-evalue)
C
Motif Enrichment Map [AGT]TAGGG[AT] (Putative HNRNPA1) (DS Set 3)

**Figure S7: Motif enrichment patterns and maps associated with additional RBPs.**
**A)** Heatmaps of minimum motif enrichment p-values across relevant junction regions for select RBP/motif pairs across DS sets tested. **B-C**) Motif enrichment maps for TIA1 (B) and HNRNPA1 (C) covering the 50 base pairs (bp) at the ends of a mis-spliced exon (orange), upstream exon (blue) or downstream exon (green), or in the 250 bp intronic regions proximal to exons.

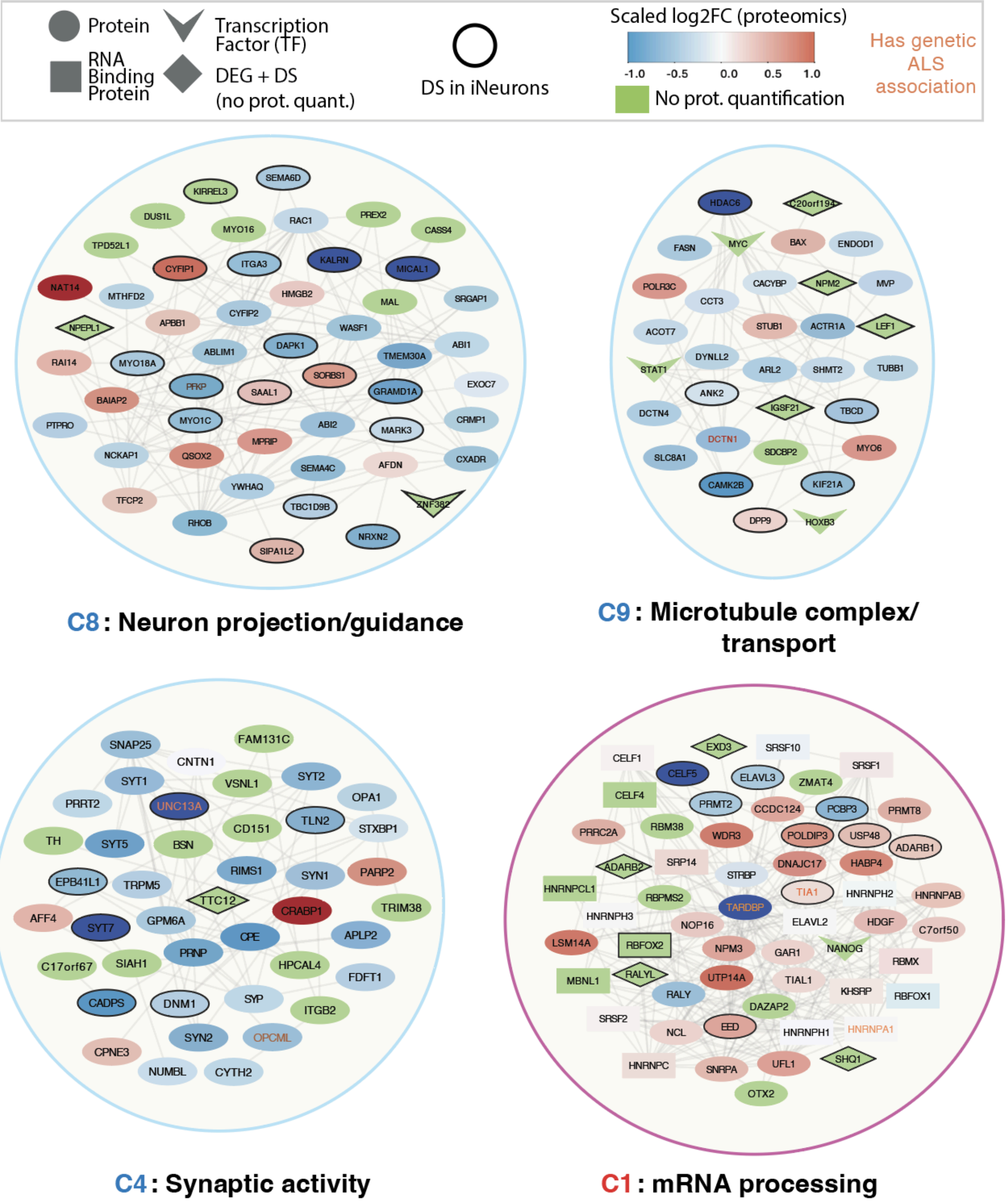


**Figure S8: Highlighted network clusters with mis-spliced genes.**
Subnetworks derived from selected clusters spanning neuronal-specific and mRNA processing pathways labeled as in Fig. 3. Featured subnetworks include proteins with DS in iNeurons (our dataset) and the union of their N nearest cluster neighborhoods, for cluster 8 (A, N = 1), cluster 9 (B, N = 1), cluster 4 (C, N = 2), and cluster 1 (D, N=1).

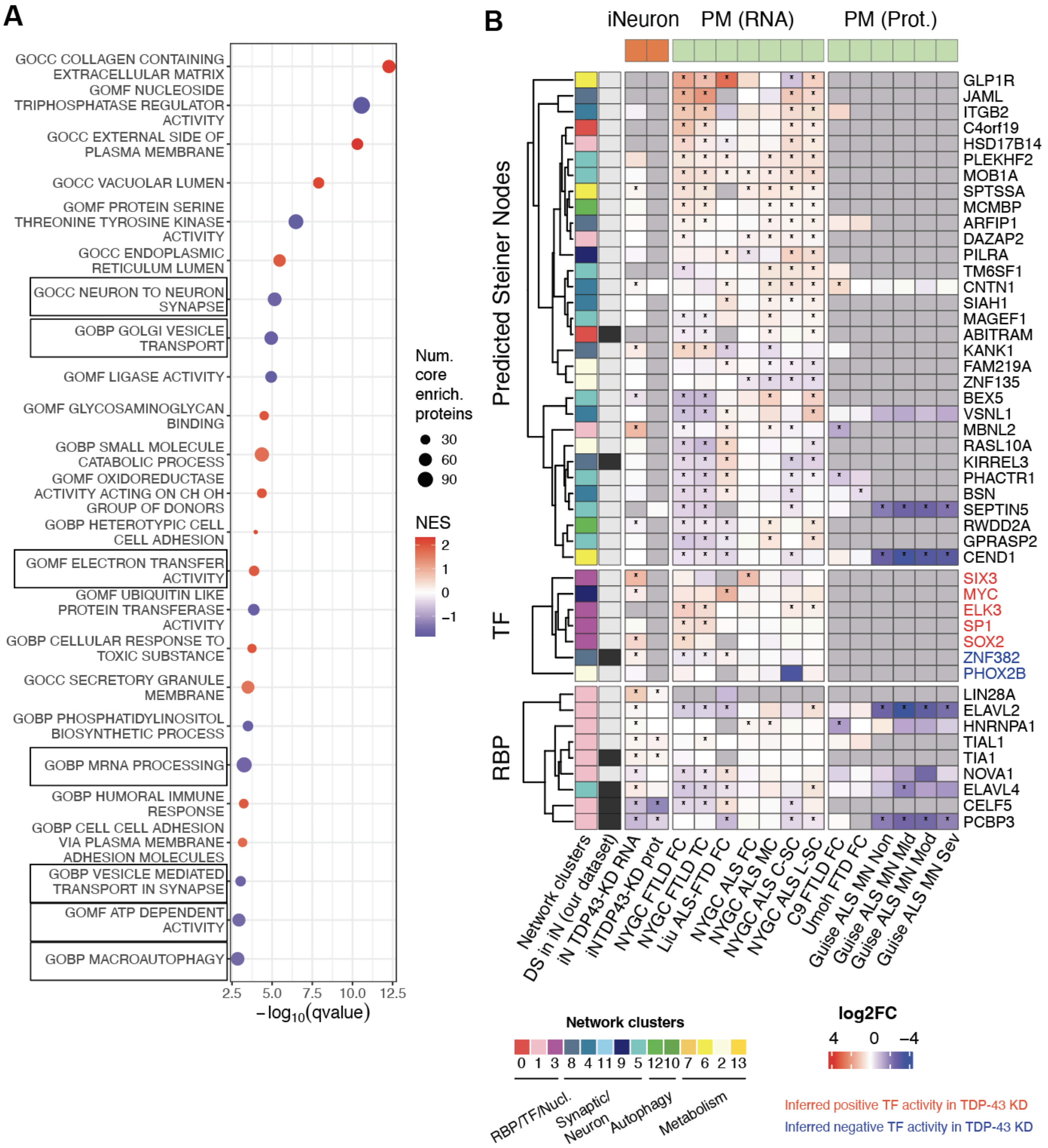


**Figure S9: Additional analyses of post-mortem datasets.**
**A)** Dotplot of GSEA results on C9-FTLD vs. control differential analysis, using MSigDB GO genesets. Normalized enrichment score (NES) indicates enrichment among ranked list of proteins relative to permuted background. Gene sets previously found to be enriched/associated with our iNeuron TDP-43 KD and network analyses are highlighted. **B**) Heatmap comparing gene and protein expression LFC values across post-mortem patient datasets for select TFs, RBPs, and predicted network Steiner nodes. * BH-adjusted p-value < 0.05.

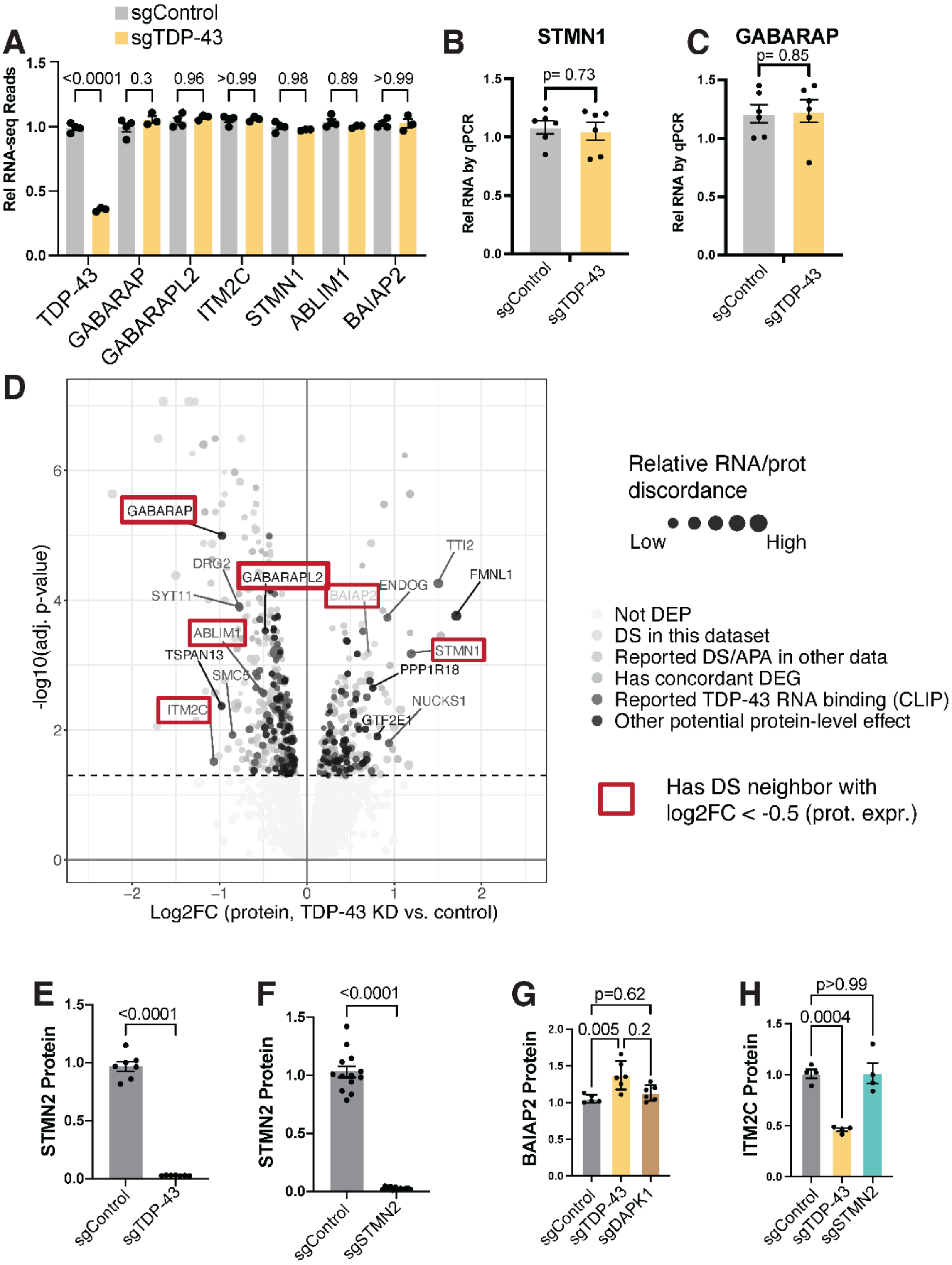


**Figure S10: Predicted functional relationships are driven by protein effects.**

**A)** Differential expression of selected genes from RNA-seq analysis shown for Control and TDP-43 KD iNeurons. p values by Ordinary one-way ANOVA with Šídák's multiple comparison test. **B)** RT-qPCR validation of STMN1 transcript levels. p values by Welch's t test; Error bars show SEM. **C)** RT-qPCR validation of GABARABP transcript levels. p values by Welch's t test; Error bars show SEM. **D)** Volcano plot of DEPs (as in Fig 1) colored by potential mechanisms contributing to observed protein expression changes. Darker grey indicates protein is less likely to be mis-spliced or regulated on a transcriptional level. Point size indicates degree of discordance between RNA and protein expression changes (larger points suggest greater discordance). **E)** Quantification of STMN2 proteins levels by western for Control KD and TDP-43 KD, Statistics using Welch's t test; Error bars show SEM. **F)** Quantification of STMN2 proteins levels by western for Control KD and STMN2 KD, Statistics using Welch's t test; Error bars show SEM. **G)** Quantification of BAIAP2 protein levels by western for Control KD, TDP-43 KD, and DAPK1 KD iNeurons for N=6 biological replicates per group. Statistics using ordinary one way ANOVA between indicated groups; Error bars show SEM. **H)** Quantification of ITM2C protein levels by western for Control KD, TDP-43 KD, and STMN2 KD iNeurons for N=4 biological replicates per group. Statistics using one way ANOVA between indicated groups; Error bars show SEM.
